# Predicting the Regenerative Potential of Retinal Ganglion Cells Based on Developmental Growth Trajectories

**DOI:** 10.1101/2025.02.28.640775

**Authors:** Joana RF Santos, Chen Li, Lien Andries, Luca Masin, Bram Nuttin, Katja Reinhard, Lieve Moons, Hermann Cuntz, Karl Farrow

## Abstract

Retinal ganglion cells in the mammalian central nervous system fail to regenerate following injury, with the capacity to survive and regrow varying by cell type. This variability may be linked to differences in developmental programs that overlap with the genetic pathways that mediate regeneration. To explore this correlation, we compared the structural changes in mouse retinal ganglion cells during development with those occurring after axonal injury. The dendritic trees of over 1,000 ganglion cells were reconstructed at different developmental stages, revealing that each cell type follows a distinct timeline. ON-sustained (sONα) cells reach maturity by P14, whereas ON-transient (tONα) cells achieve their maximum dendritic size by P10. Modeling of the dendritic changes indicate that while sONα and tONα follow similar growth programs the onset of growth was later in sONα. After optic nerve crush, the remodeling of dendritic architecture differed between the two cell-types. sONα cells exhibited rapid dendritic shrinkage, while tONα cells shrank more gradually with changes in branching features. Following injury, sONα cells reverted to an earlier developmental state than tONα cells. In addition, after co-deletion of PTEN and SOC3, neurons appeared to regress further back in developmental time. Our results provide evidence that a ganglion cell’s resilience to injury and regenerative potential is predicted by its maturation timeline. Understanding these intrinsic differences could inform targeted neuroprotective interventions.

## Introduction

In the adult mammalian central nervous system (CNS), axonal injuries typically result in permanent damage, as most neurons fail to regenerate their lost axons or restore synaptic connections (1). While neurons exhibit intrinsic regenerative potential during development, as they extend axons toward target brain regions, this ability declines rapidly within the first weeks of life, as the CNS transitions from a state of dynamic growth to structural stability (2,3).

Analysis of the molecular programs involved reveals that while the capacity for axonal repair diminishes with maturation, neuronal regeneration, and development rely on shared molecular pathways (4). Given this connection, we hypothesized that examining dendritic growth trajectories during development is likely to provide insight into the mechanisms governing neuronal survival and regenerative potential after injury. Single-cell transcriptomic studies using the optic nerve crush (ONC) model, which causes a loss of ~80% of ganglion cells (5), reveal striking variability in survival and regenerative potential across the ~40 ganglion cell subtypes, with survival rates ranging from 1% to 98% (6). Among these subtypes, ON-sustained (sONα) and ON-transient (tONα) alpha ganglion cells exhibit distinct regenerative capacities, despite their transcriptional similarity. sONα cells demonstrate significantly higher resilience to injury than tONα cells (6). Axonal regeneration following ONC is typically absent, but targeted manipulations have shown limited success. In particular, deleting PTEN and SOCS3, key inhibitors of the JAK/STAT3 and PI3K/mTOR pathways, enhances axonal regrowth (7,8). While PTEN deletion alone predominantly promotes regeneration in alpha ganglion cells, combining it with CNTF and SOCS3 deletion amplifies survival and regeneration across a broader spectrum of retinal ganglion cells. Similarly, developmental genes such as Stat3, Sox11, and Tubb3 also promote axon regrowth, highlighting the shared molecular mechanisms between development and injury recovery (9).

It has been observed that interactions between different neuronal compartments play an important role both during development and during regeneration after injury. During development axons tend to innervate their target before dendrites mature and develop, suggesting a clear order of events (10–13). A similar order of events has been suggested to be useful for post-injury regeneration. In Caenorhabditis elegans, dendrites appear to actively suppress ectopic sprouting, as acute lesioning of dendrites tends to promote axon regeneration after injury (14). A similar antagonistic interaction is observed in zebrafish, where the dendrites of ganglion cells shrink following axonal injury, suggesting that dendritic shrinkage may facilitate axon regrowth (15). These findings collectively highlight how dendritic across the retina, indicating no sampling bias. The orientation is nasal (N) to temporal (T). remodeling may play a pivotal role in facilitating axonal regeneration, underscoring the intricate interplay between neuronal compartments during recovery.

In this study, we systematically compare the dendritic morphology in ganglion cells during development and after optic nerve injury, both with and without PTEN/SOCS3 deletion. We focus on sONα and tONα cells, which exhibit distinct resilience to injury (6). These two cell-types are of particular interest as they are orthologues of parasol and midget ganglion cells in the human retina (16). Our analysis revealed three key findings: first, the timing of developmental growth varies between cell types; second, both cell-types undergo shrinkage after injury and during axonal regeneration; third, modeling their dendritic growth and shrinkage patterns showed that these injured morphologies correspond to different developmental ages, suggesting that developmental timing is related to a cell-types resilience and regenerative potential.

## Results

### Characterizing the anatomy of retinal ganglion cells in the mouse retina

To examine how neuronal development influences their response to injury, we analyzed the dendritic morphology in Thy1-YFP-H mice across multiple developmental stages and post-injury time points (17–19) (Figure 1A). We first characterized the dendritic structures of healthy adult ganglion cells, measuring dendritic depth relative to starburst amacrine cells labeled with choline acetyltransferase (ChAT; magenta; Figure 1B). Alpha ganglion cells were identified using an antibody against neurofilament protein (20–24) (SMI-32; red; Figure 1C). The dendritic trees of individual ganglion cells were traced using a semi-automated algorithm (Figure 1D and E) (25). Each cell was categorized using a combination of its dendritic tree depth and SMI-32 labeling. Cells were classified into one of the four alpha ganglion cell types: ON-sustained alpha (sONα), ON-transient alpha (tONα), OFF-sustained alpha (sOFFα), OFF-transient alpha (tONα) or other (Figure 1F).

**Figure 1.**
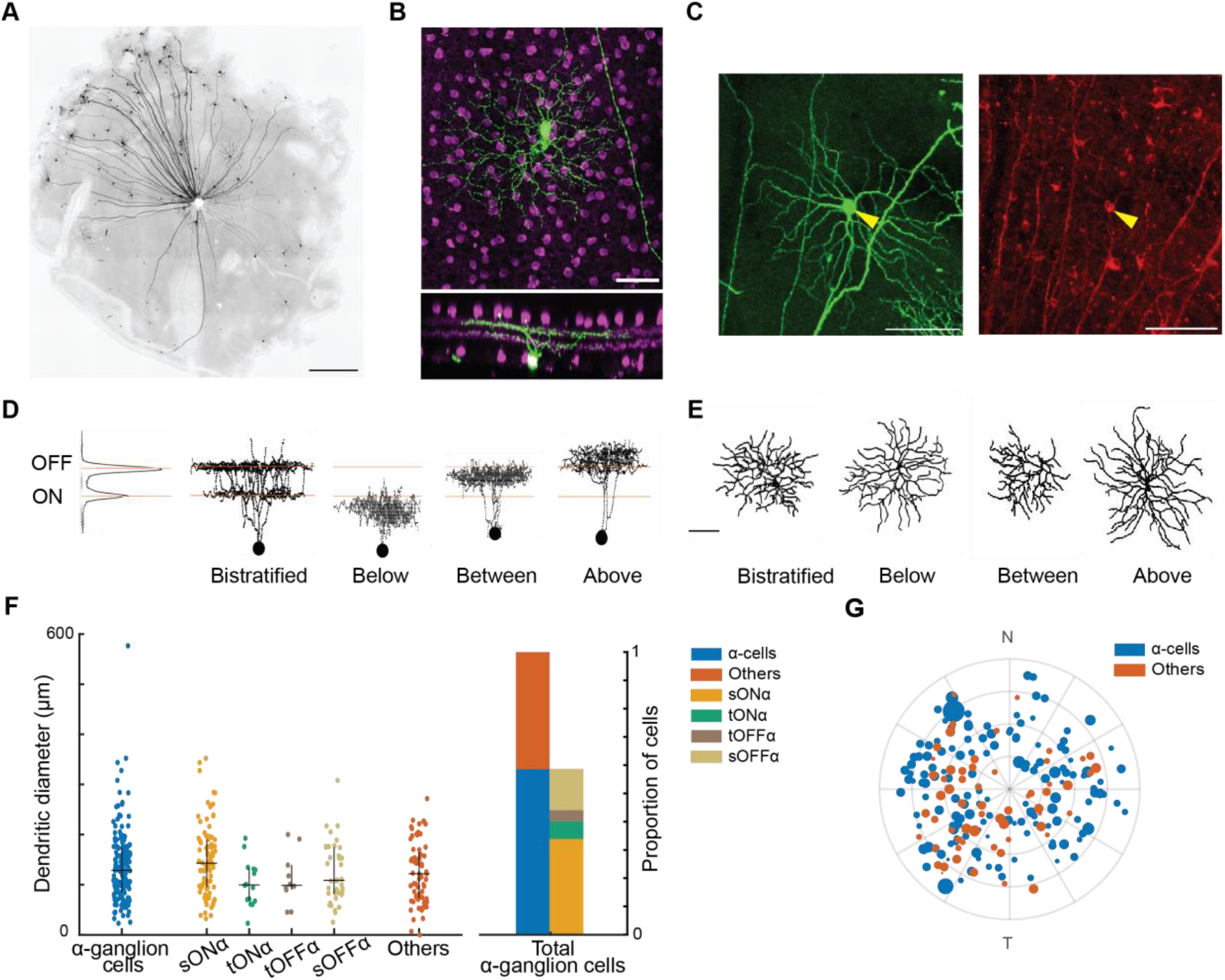
Classification of ganglion cells based on dendritic morphology and stratification. **(A)** Whole-mount retina from a Thy1-YFP-H mouse, showing the characteristic sparse labeling of ganglion cells in this model. Scale bar: 100 µm. (B)Example of a labeled ganglion cell. Top: en-face view of the dendritic arbor. Bottom: side-view, showing dendritic stratification within the inner plexiform layer (IPL). The green channel represents the labeled ganglion cell, while the magenta channel marks starburst amacrine cells expressing choline acetyltransferase (ChAT). Scale bar: 20 μm. **(C)** Identification of alpha ganglion cells using SMI32 immunolabeling. Left: a YFP-labeled ganglion cell (green). Right: the same cell labeled with SMI32 (red), indicating neurofilament expression characteristic of alpha ganglion cells. Scale bar: 20 μm. **(D-E)** Representative examples of ganglion cells stratifying at different depths within the IPL. Left: side-view, showing the relationship of the dendrites to the ChAT bands (orange lines). Cells are grouped into four stratification patterns: bistratified (first column), below the ChAT bands (second column), between the ChAT bands (third column), and above the ChAT bands (fourth column). Right: respective en-face view of the dendritic arbor. Scale bar: 20 μm. **(F)** Thy1-YFP-H mice preferentially label alpha ganglion cells. Left: Dendritic diameter of alpha and non-alpha ganglion cells in adult mice. Black lines indicate the median and interquartile range (IQR; 25th and 75th percentiles). Right: Distribution of alpha and non-alpha ganglion cells, showing that alpha cells constitute ~60% of the labeled population, with a bias toward ON alpha subtypes. **(G)** Spatial distribution of analyzed ganglion cells. Alpha ganglion cells (dark blue) and non-alpha ganglion cells (dark orange) were evenly distributed across the retina, indicating no sampling bias. The orientation is nasal (N) to temporal (T).

Analysis of the Thy1-YFP-H mouse line revealed an overrepresentation of alpha ganglion cells compared to their actual proportion (~5%) in the total ganglion cell population (26). Among the 266 ganglion cells traced (n = 266 cells, 12 retinas, 7 Santos et al., Sep 2025 – preprint copy – BioRxiv mice aged 6-8 weeks), 67% (179/266) were identified as alpha ganglion cells. Additionally, ~10% were bistratified (27/266), and 5% (12/266) expressed CART, a marker for direction-selective ganglion cells. The spatial distribution of all ganglion cells is reported in Figure 1G and Supplementary Figure 1. Our subsequent analysis focused on sONα and tONα ganglion cells which are preferentially labeled in the Thy1-YFP-H mouse line. These two cell types have been shown to differ in their resilience to axonal injury and specific regenerative treatments, making them relevant for further investigation (6,27–29).

### Ganglion cells reach mature length at eye-opening

To determine the timeline of ganglion cell maturation, we analyzed dendritic morphology across multiple postnatal stages, tracking changes in dendritic growth and branching features. We reconstructed the dendritic architecture of 1,368 retinal ganglion cells across five developmental time points (P03, P07, P10, P14, and P28) to quantify growth patterns (Figure 2A-B). We characterized each cell’s dendritic tree using 11 morphological parameters (Figure 2C), which included metrics describing the size of the dendritic (e.g., total dendritic length, total surface area, Sholl analysis index, mean Euclidean distance and mean path length) represented on the left of the star plots), and metrics that capture its branching features (e.g., number of branches, mean branch order, mean branching angle, centripetal bias, convexity and balancing factor) represented on the right side of the star plots (25,30). Additionally, convexity and area of the soma were included as measures of shape. A detailed description of these parameters can be found in the Methods section and Supplementary Figure 2.

**Figure 2.**
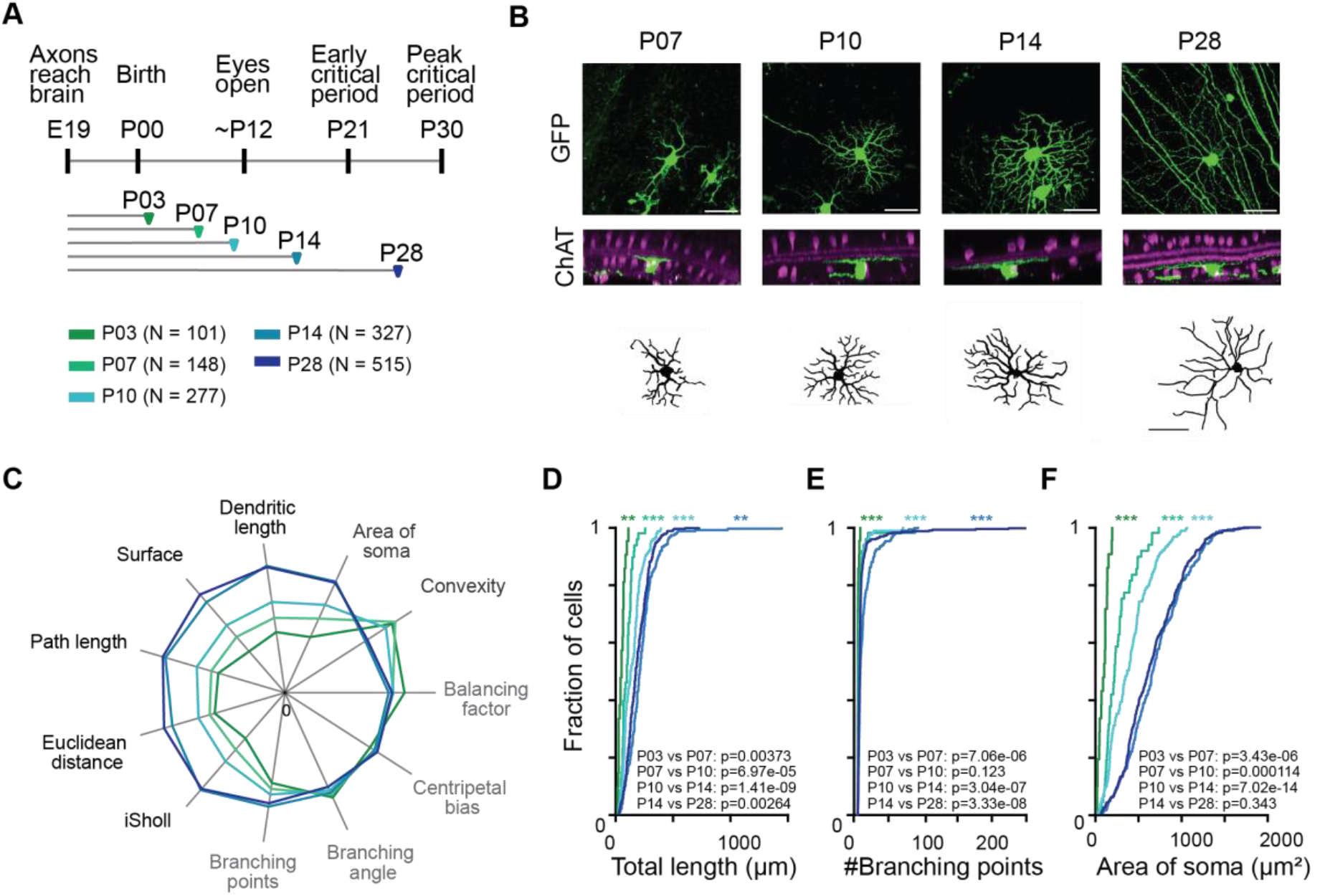
Ganglion cell dendrites grow until eye-opening and stabilize thereafter. **(A)** Schematic of developmental time points analyzed: postnatal day (P) 3, P7, P10, P14, and P28). **(B)** Representative ganglion cells at each stage. Top: En-face confocal images showing dendritic morphology (green). Middle: Side-view confocal images of the same cells, with ChAT-positive starburst amacrine cells in magenta. Bottom: Traced en-face reconstructions of the corresponding dendritic arbors. Scale bar: 20 µm. **(C)** Quantification of dendritic morphology across development using 11 structural parameters. Left: size-related parameters including total dendritic length, dendritic surface area, mean path length, and mean Euclidean length (black). Right: branching features including branching number, mean branch order, mean branching angle, convexity, and balancing factor (light gray). Shape parameters like soma size and convexity are represented in dark gray. **(D - F)** Cumulative distribution plots for dendritic length, branching points, and soma area across developmental stages. Statistical comparisons between groups were performed using a Kolmogorov-Smirnov (KS) test, with significant shifts in distribution observed across stages (marked with (*, **, ***) representing *p* ≤ 0.05, *p* ≤ 0.01, or *p* ≤ 0.001, respectively).

We identified three distinct phases of dendritic growth in retinal ganglion cells. First, an expansion phase (P03–P14), during which dendritic size increased significantly. Dendritic length peaked at 1280.84 µm by P14 (p = 1.30 × 10^−22^), with the most pronounced growth occurring between P10 and P14 (p = 2.55 × 10^−9^). Dendritic surface area followed a similar trajectory, increasing from 29,504.63 µm^2^ (P03) to 108,685.85 µm^2^ (P14) (p = 5.43 × 10^−14^). Soma size also increased, from 87.57 µm^2^ (P03) to 608.57 µm^2^ (P14) (p = 4.17 × 10^−18^), reflecting active dendritic and somatic expansion. Second, a structural refinement phase (P10– P14), during which dendritic organization was further modified. Branching points remained stable before P10 but increased significantly between P10 and P14 (p = 3.09 × 10^−5^). Convexity decreased slightly from P07 to P10 (p = 3.19 × 10^−3^) before increasing sharply between P10 and P14 (p = 4.16 × 10^−12^), then stabilizing. Branching features such as mean branching angle remained unchanged, indicating that dendritic refinement was limited to select structural features.

Finally, a stabilization phase (P14–P28), in which dendritic length decreased slightly (p = 3.11 × 10?4). Dendritic surface area, Euclidean distance, and path length did not change significantly. Dendritic branching features remained stable, except for minor refinements in branching angle (p = 2.60 × 10?4), confirming that dendritic architecture was largely set by P14.

These results define a three-phase model of dendritic maturation, with rapid expansion until P14, selective refinement from P10 to P14, and stabilization beyond P14, delineating the structural constraints of retinal ganglion cell development.

### sONα and tONα alpha ganglions mature at different times

To assess differences in dendritic development across ganglion cell types, we analyzed sONα and tONα cells at four postnatal stages: P07, P10, P14, and P28. P03 was excluded because ChAT bands had not yet matured, making cell type classification unreliable (31). sONα and tONα cells exhibit distinct growth trajectories (Figure 3A-B; Supplementary Figure 3). sONα cells increased significantly between P07 and P10 (+46.89%, p = 0.002, KS test), followed by further growth from P10 to P14 (+104.96%, p = 4.79 × 10 ^-11^, KS test) before stabilizing by P28 (Figure 3C-D; Supplementary Figures 4–5). In contrast, tONα cells did not show significant changes in dendritic length between P07 and P10 (p = 0.152, Mann-Whitney, ANOVA: p = 0.018) or P10 and P14 (p = 0.004, Mann-Whitney), yet exhibited a significant overall increase from P07 to P28 (p = 0.001, Mann-Whitney). This pattern coincided with a significant increase in the number of branching points in tONα cells between P07 and P10 (p = 0.016, ANOVA) and P10 and P14 (p = 0.013, ANOVA), suggesting that while major dendritic expansion slowed, arborization continued through a gradual addition of new branches (Figure 3C). Despite differences in growth rates, branching features remained stable in both cell types across development. The dendritic shape evolved differently. Convexity, a measure of dendritic shape, increased significantly in sONα from P10 to P14 (p = 0.003, ANOVA), while remaining unchanged in tONα (p = 0.416, ANOVA) (Figure 3C). Branching angles did not change significantly in either subtype, confirming that increases in dendritic size were not accompanied by major shifts in branch organization (Figure 3D).

**Figure 3.**
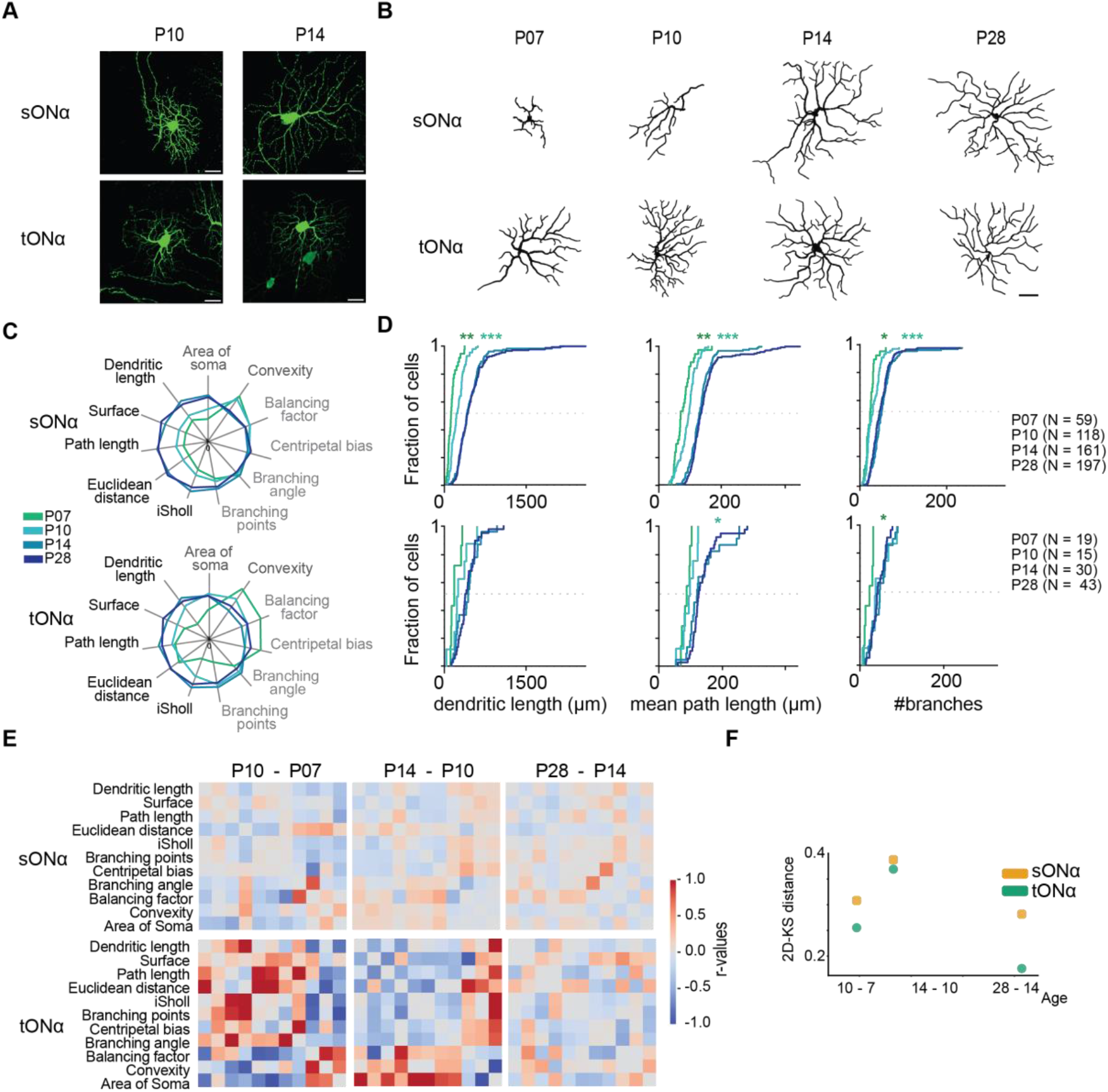
sONα and tONα alpha ganglion cells follow distinct developmental growth profiles. **(A)** Representative confocal images of sONα (top) and tONα (bottom) ganglion cells at different developmental stages (P7, P10, P14, P28). Scale bars: 50 µm. **(B)** Traced dendritic arbors of sONα (top) and tONα (bottom) cells, illustrating differences in growth patterns across postnatal stages. **(C)** Quantification of dendritic morphology over the development of sONα (top) and tONα (bottom). Left: size-related parameters (black). Right: branching features (light gray). **(D)** Cumulative growth curves of total dendritic length, mean path length and branch number for sONα (top) and tONα (bottom), tracking their developmental progression. Statistical comparisons between groups were performed using a KS-test or Mann-Whitney U test (Wilcoxon rank-sum test) based on sample size (n < 20), with significant shifts in distribution observed across stages (marked with (*, **, ***)). **(E)** Differential correlation heatmaps showing structural changes between successive developmental stages. Pearson correlation matrices from each stage are subtracted from the preceding stage to visualize shifts in dendritic architecture over time. Gray regions indicate minimal change, while colored regions highlight morphological differences between time points. Top: sONα cells; Bottom: tONα cells. **(F)** KS distances quantifying structural changes in dendritic morphology across developmental time points. The 2D KS distance compares the distribution of upper-triangle correlation coefficients between successive stages, measuring the magnitude of structural reorganization. Data points represent individual replicates (orange circles: sONα; green squares: tONα); larger KS distances indicate greater morphological shifts. Data were collected from 56 retinas, with 7 mice per time point.

To assess dendritic stability over time, we computed correlation matrices comparing dendritic parameters across developmental stages (Figure 3E). sONα cells exhibited high temporal stability (P07–P10: r = 0.90; P10–P14: r = 0.96; P14–P28: r = 0.96), while tONα cells showed greater early variability (P07–P10: r = 0.56; P10–P14: r = 0.69), stabilizing by P14–P28 (r = 0.90). We next quantified developmental shifts using the Kolmogorov-Smirnov (KS) distances (Figure 3F). KS distances were consistently larger in tONα cells, indicating greater morphological reorganization between stages. Differences were significant before P14 (P07– P10: p = 0.0011; P10–P14: p = 0.0003) but not thereafter (*p* = 0.54), suggesting that both cell types converge in mature morphology.

Thus, tONα cells undergo rapid early expansion before stabilizing at P10, while sONα cells develop more gradually, continuing growth until P14. These intrinsic differences in maturation may influence how each cell type responds to injury.

### sONα and tONα alpha ganglions respond differently to injury

Next, we examined the effects of injury on the dendritic structure of sONα and tONα ganglion cells (Figure 4A; Supplementary Figure 6). We subjected Thy1-YFP-H mice to an optic nerve crush, six to eight weeks after birth. Retinas were collected from uninjured mice and from animals at five different time points after injury: 0, 1, 4, 7, and 14 days post-injury (dpi). A total of 938 alpha ganglion cells were examined (41 retinas, 5– 6 mice per time point).

**Figure 4.**
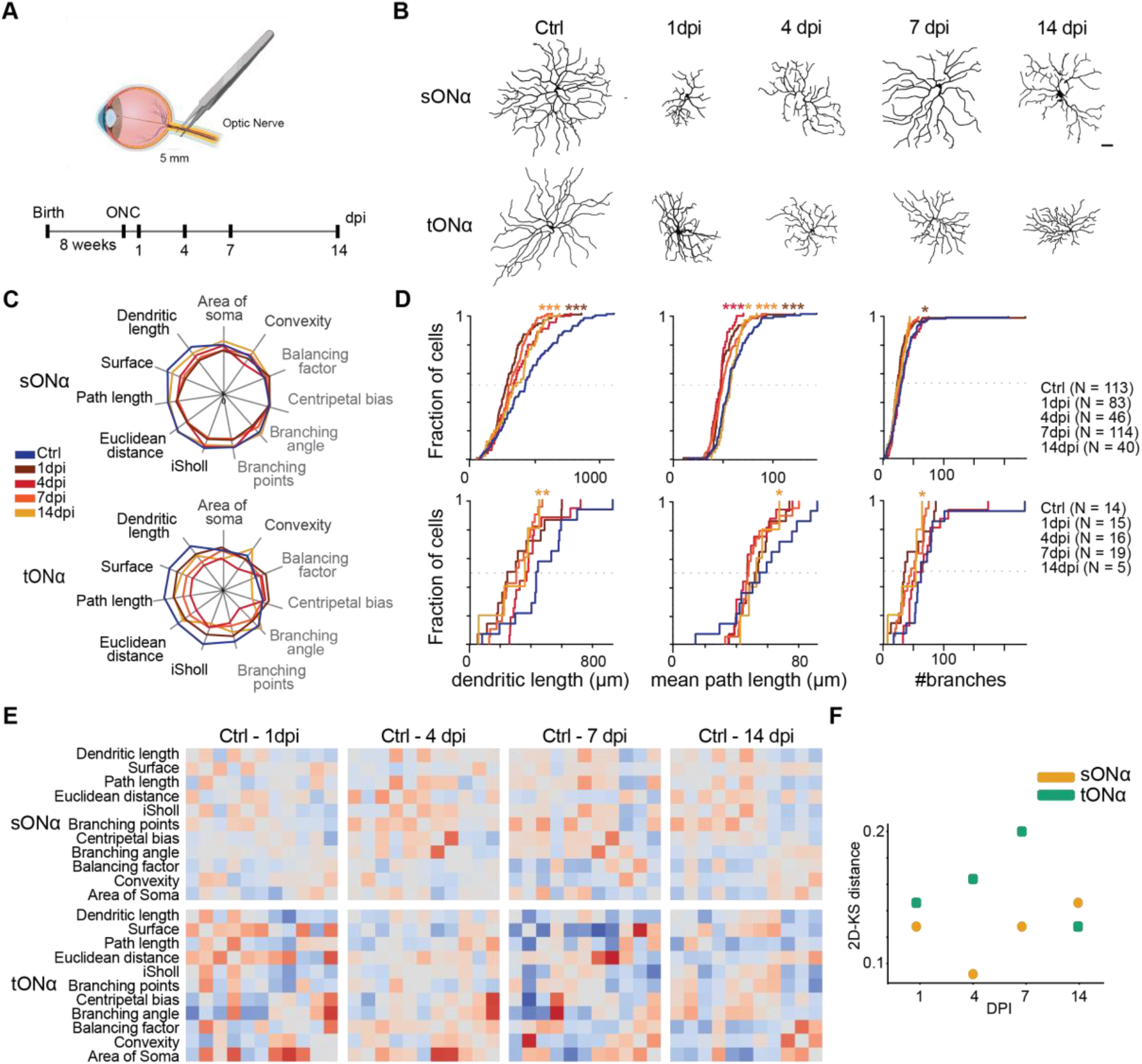
sONα and tONα ganglion cells exhibit distinct dendritic remodeling patterns following injury. (A)Schematic representation of the optic nerve crush (ONC) procedure, with the timeline indicating uninjured (Ctrl) and post-injury analysis points (1, 4, 7, and 14 days post-injury, dpi). **(B)** Representative illustrations of sONα and tONα cell dendritic morphologies in uninjured and injured conditions, highlighting progressive structural changes in each cell-type. Scale bar = 50 µm. **(C)** Quantification of dendritic morphology at different post-injury stages for sONα (top) and tONα (bottom). Left: size-related parameters (black). Right: branching features (light gray). **(D)** Cumulative growth curves of total dendritic length, mean path length, and branch number for sONα (top) and tONα (bottom), tracking their developmental progression. Statistical comparisons between groups were performed using a KS-test or Mann-Whitney U test (Wilcoxon rank-sum test) based on sample size (n < 20), with significant shifts in distribution observed across stages (marked with (*, **, ***)). **(E)** Differential correlation heatmaps showing shifts in dendritic morphology over time. Pearson correlation matrices at each post-injury stage are subtracted from the baseline (0 dpi) correlation matrix to highlight structural deviations. Gray regions indicate minimal change, while colored regions reflect stronger deviations from the baseline. Top: sONα cells; Bottom: tONα cells. **(F)** KS distance quantifying morphological deviations from baseline (0 dpi). Data points represent individual replicates (orange circles: sONα; green squares: tONα); larger KS distances indicate greater deviations from baseline. Data were collected from 41 retinas, with 5–6 mice per time point.

sONα and tONα ganglion cells exhibited distinct dendritic remodeling patterns following injury (Figure 4A; Supplementary Figure 6). Both cell types retracted significantly, but the timing and extent of regression differed. sONα cells shrank immediately, with dendritic length reduced from 1180.4 µm to 675.7 µm by 1 dpi. In contrast, tONα cells remained stable at 1 dpi but showed a significant reduction by 7 dpi (Mann-Whitney, p = 0.007). By 14 dpi, this difference was no longer significant (p = 0.09), suggesting stabilization (Figure 4B-D; Supplementary Figures 7– 8). Dendritic branching features evolved differently between cell types. sONα cells lost branches early (ANOVA, p = 0.047), while tONα cells maintained branch number (ANOVA, p = 0.30). Centripetal bias increased in sONα (KS test, p = 0.001), indicating retraction toward the soma, whereas tONα showed no significant shift (ANOVA, p = 0.41). Branching angles also diverged: sONα exhibited a branching reorganization post-injury (KS test, p = 0.0195; ANOVA, p = 0.01), whereas tONα angles remained stable (ANOVA, p = 0.3045). The dendritic shape was also affected.

Morphological stability over time was assessed using correlation matrices comparing dendritic parameters across post-injury stages to the baseline (0 dpi) (Figure 4E). At 1 dpi, sONα cells showed only minor deviations from baseline, whereas tONα cells exhibited larger structural shifts. By 14 dpi, sONα cells remained largely unchanged, while tONα cells continued to deviate from their baseline morphology, indicating persistent instability of their dendritic organization. The Kolmogorov-Smirnov (KS) distance quantified the extent of morphological changes (Figure 4F). While both cell types deviated from baseline post-injury, tONα cells exhibited larger shifts in correlation structure, reflecting greater instability. However, direct comparisons of sONα and tONα correlation values did not reach statistical significance (Mann-Whitney U test; 1 dpi: p = 0.09; 4 dpi: p = 0.40; 7 dpi: p = 0.55; 14 dpi: p = 0.94).

These findings indicate that both sONα and tONα cells undergo substantial dendritic shrinkage after axonal injury. However, sONα cells respond immediately and stabilize more quickly, whereas tONα cells display delayed and more variable remodeling.

### sONα and tONα ganglion cells retain their function

To determine whether structural changes affected neuronal function, we performed patch-clamp recordings of sONα and tONα ganglion cells at different time points following optic nerve crush (Figure 5A-B, Supplementary Figure 9A). A total of 157 ganglion cells were recorded from 10 retinas: 50 cells at 1 dpi, 58 at 4 dpi, and 49 at 14 dpi. Cells were classified as sONα or tONα based on their responses to a spot stimulus varying at different sizes and response to a chirp stimulus(32) (Figure 5A-B, Supplementary 9A). For a subset of patched cells, morphology was confirmed post hoc. The analysis of functional properties focused on these identified alpha cells.

**Figure 5.**
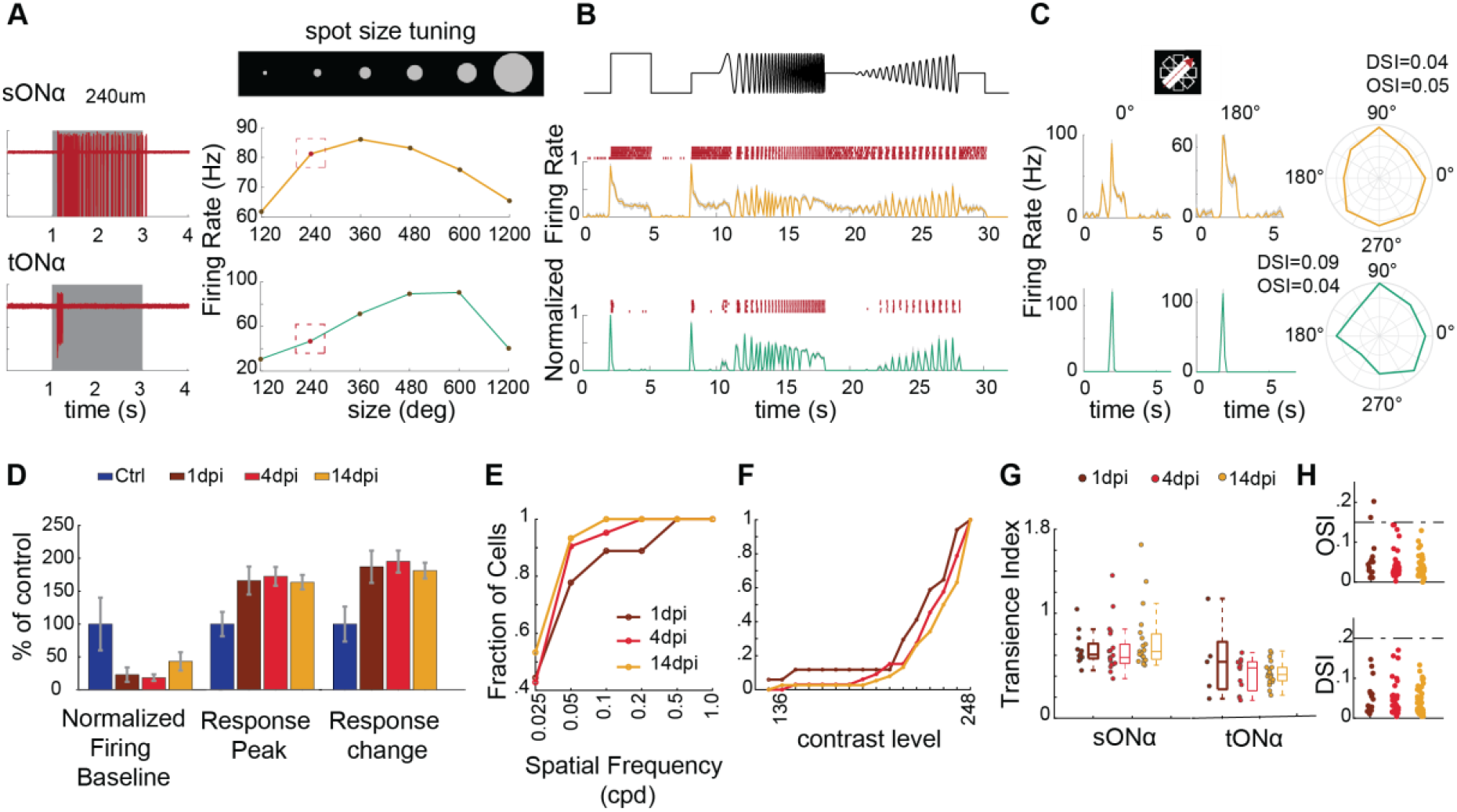
Functional properties of retinal ganglion cells following optic nerve crush injury. **(A)** Example responses of sONα (top) and tONα (bottom) ganglion cells to a 240 µm spot stimulus presented on a black background for 2 seconds. Left: Raster plots showing spike times over repeated trials (red). Right: Firing rate curves (128 ms bins) across trials. Gray bars indicate stimulus duration. **(B)** Responses of the same example cells to a chirp stimulus, which consists of a slow contrast modulation followed by a frequency modulation designed to assess temporal processing. Top: Raster plots showing individual spike events (red). Bottom: Mean firing rates across trials (colored traces). **(C)** Direction tuning properties of ganglion cells in response to a moving bar stimulus. Left: Example response traces to bars moving at 0° and 180°. Right: Polar plot illustrating directional tuning for the same cells. **(D)** Quantification of firing rates in patched ON alpha ganglion cells. Bar plots show baseline firing rate before stimulus onset (left), peak response to a moving bar (middle), and relative change in response compared to baseline (right), separated by condition (Ctrl, 1, 4, and 14 dpi). **(E)** Cumulative distributions of preferred spatial frequencies across all recorded ON alpha ganglion cells at different dpi in response to drifting gratings. **(F)** Cumulative distributions of preferred contrast levels across all recorded ON alpha ganglion cells at different dpi. **(G)** Transience index of sONα (left) and tONα (right) at different dpi, measuring the stability of transient responses over time. **(H)** Direction selectivity index (DSI, top) and orientation selectivity index (OSI, bottom) at different dpi, showing that selectivity remains stable after injury. Data were collected from 157 ganglion cells recorded from 10 retinas (50 cells at 1 dpi, 58 at 4 dpi, 49 at 14 dpi).

Baseline firing rates decreased significantly across all post-injury time points compared to controls but partially recovered by 14 dpi (Figure 5D, Supplementary Figure 9E). The firing baseline was measured during the 2 seconds before the onset of a moving bar stimulus. We also analyzed the peak response during the stimulus and found increased response amplitudes, along with changes in relative response changes. To assess spatial processing, we presented drifting gratings at different spatial frequencies. At 1 dpi, most cells responded to a broader range of spatial frequencies compared to later time points (Figure 5E).

Analysis of responses to a contrast ramp in the ‘chirp’ stimulus revealed strong responses to high contrast across all post-injury time points. While differences between 1, 4, and 14 dpi were subtle, cells at 1 dpi tended to show greater sensitivity, with half responding at lower contrast levels. The transience index, calculated from the area under the post-stimulus time histogram (PSTH), showed similar distributions across all time points. Direction selectivity (DSI), orientation selectivity (OSI), and ON-OFF indices also remained stable from 1 to 14 dpi (Figure 5H, Supplementary Figure 9F).

These findings indicate that despite structural remodeling, sONα and tONα ganglion cells retain core aspects of their functional identity post-injury despite dendritic shrinkage. This persistence of functional responses suggests that at least some synaptic connections remain intact, indicating that the cells continue to receive and process signals after injury.

### Comparison of Development and Response to Injury in alpha retinal ganglion cells

To assess parallels between normal development and the response to injury, we compared the dendritic structure of sONα and tONα ganglion cells at postnatal stages (P07, P10, P14, P28) with their morphology at different post-injury time points (1, 4, 7, and 14 dpi). Previously, we found that dendritic growth occurs primarily between P07 and P14, stabilizing by P28. After injury, the most rapid dendritic shrinkage occurs at 1 dpi, with partial recovery by 4 dpi and stabilization at later time points. sONα cells show a transient loss followed by partial restoration, while tONα cells exhibit a more gradual decline with little recovery.

Correlation analyses between development and injury revealed that sONα cells largely retained their developmental relationships, except at 1 dpi, where this correlation was disrupted. By 4 dpi, their morphology realigned with P14-P28 stages. In contrast, tONα cells showed little correlation between injury and development across all time points, indicating a distinct remodeling trajectory (Figure 6A-B). Examining specific parameters, both cell types exhibited an initial regression in dendritic length to P10-like values following injury (Figure 6C). However, while sONα cells stabilized at this stage, tONα cells continued to regress, reaching P07-like values by 4 dpi. In sONα cells, dendritic shrinkage occurred without major changes in branching, and branch organization remained largely intact. In contrast, tONα cells showed a more gradual reduction in dendritic length, but branching measures remained stable across time points (Figure 6C).

**Figure 6.**
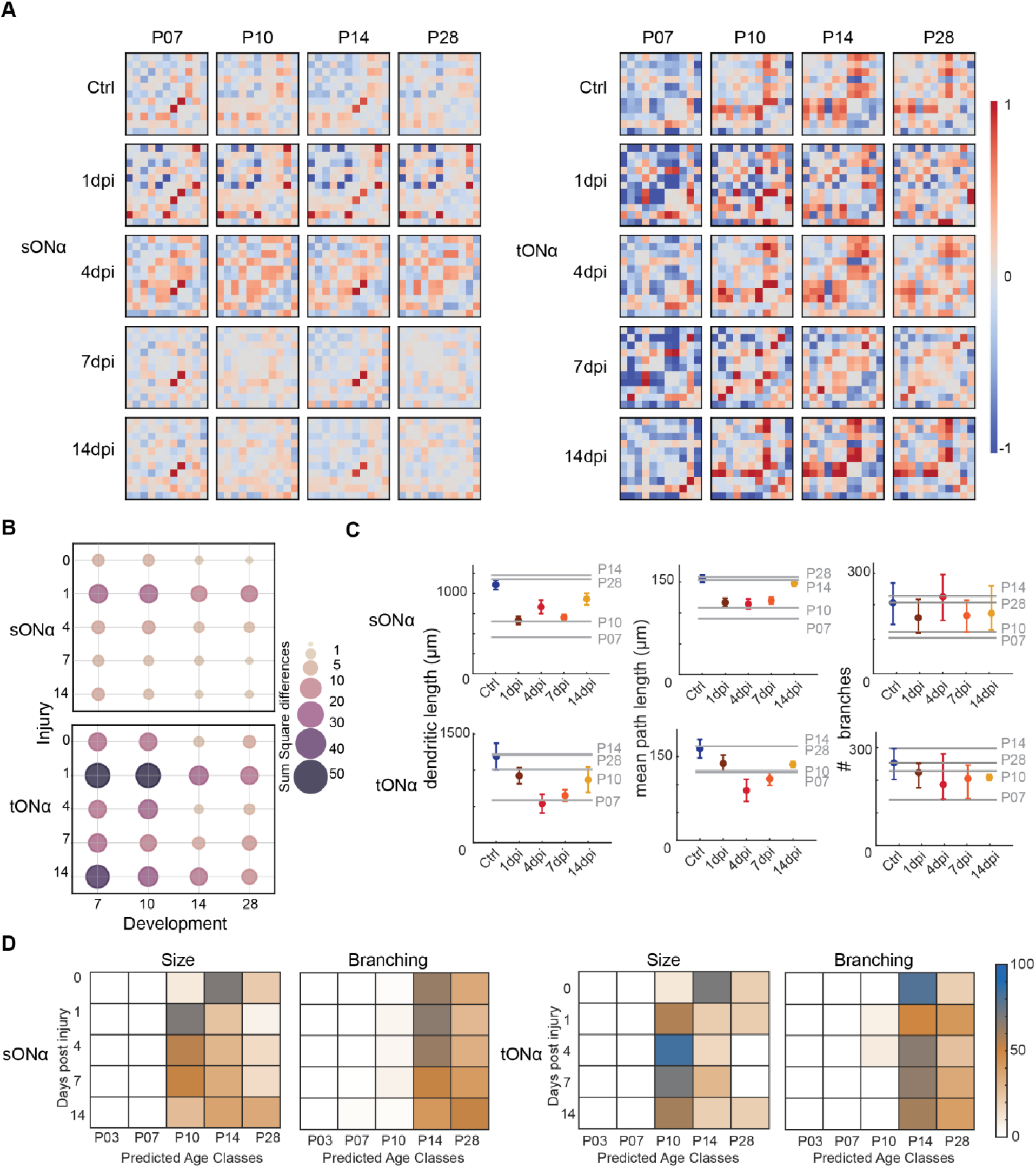
Comparison of developmental and injury-induced dendritic remodeling in ganglion cells. **(A)** Differential Pearson correlation heatmaps comparing developmental and injury conditions in sONα (left) and tONα (right) ganglion cells. Each heatmap represents the subtraction of a correlation matrix from a developmental time point (P03, P07, P10, P14, and P28 - horizontal axis) and an injury time point (control, 1, 4, 7, and 14 dpis - vertical axis). Warmer colors indicate greater correlation differences, with 0 representing no change. **(B)** Bubble plot summarizing the sum of squared differences from the heatmaps in (A) in sONα (top) and tONα (bottom). The color scale ranges from beige to dark purple, representing the magnitude of the squared differences. **(C)** Error plots of dendritic length, path length, and number of branches across post-injury stages. Colored error bars represent the mean and standard error of the mean (SEM) for each post-injury timepoint, while gray horizontal lines indicate the median values at corresponding developmental stages. **(D)** Confusion matrices displaying k-nearest neighbors (k-NN) classifier results for predicting the developmental equivalence of injured ganglion cells. The classifier was trained separately for sONα and tONα cells using either size-related metrics (left matrix) or branching-related metrics (right matrix). Rows represent the true injury timepoints, while columns indicate the predicted developmental stages. The color intensity reflects the proportion of predictions, with higher values indicating a stronger classification match.

A k-nearest neighbors (k-NN) classifier, trained separately on size and branching features, was used to compare injured ganglion cells to developmental stages (Supplementary Figure 10). When using size-related parameters such as total dendritic length and surface area, the classifier assigned early post-injury stages (1 dpi and 4 dpi) to P10-like stages. As recovery progressed, the predicted developmental stage increased, with later time points aligning more closely with P14 and P28 (Figure 6D). However, when classifying based on branching-related features such as branch order and density, injured cells were consistently assigned to later developmental stages (P14-P28) at all post-injury time points. This suggests that while dendritic size regresses to earlier developmental states after injury, branching remains largely unaffected in sONα cells, but tONα cells deviate from normal development (Figure 6D).

### Modeling cell-type-specific dendritic growth during development

To investigate whether dendritic remodeling after injury mirrors developmental growth patterns, we employed a computational model to simulate dendritic expansion in sONα and tONα ganglion cells. This model reconstructs synthetic dendritic trees by optimizing branch placement and spatial coverage under biologically relevant constraints (33) (Figure 7A; Supplementary Figure 11). By iterating through growth parameters, the model allows us to compare natural developmental trajectories with post-injury remodeling, revealing potential predictive relationships between these processes.

**Figure 7.**
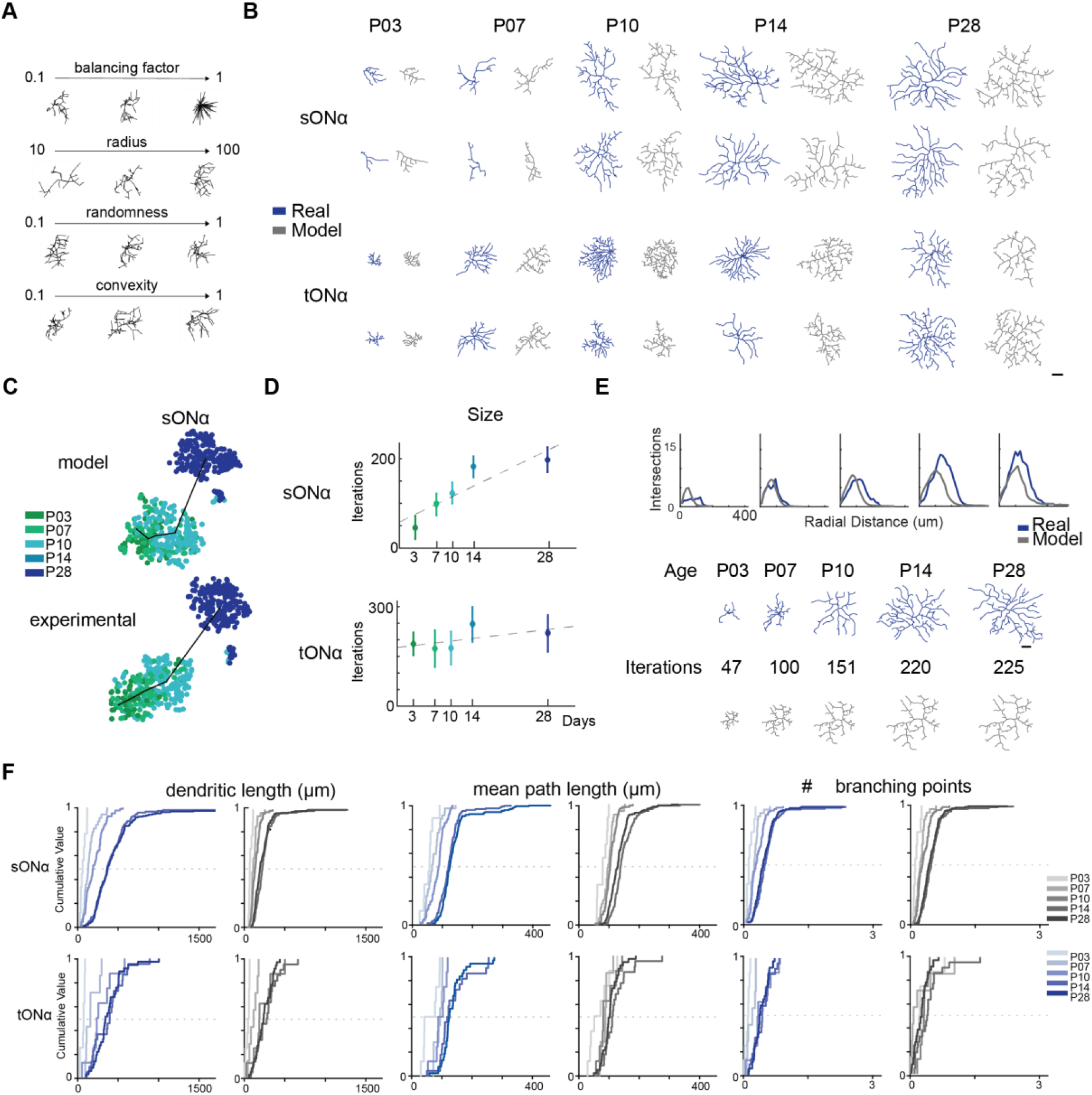
Computational modeling of dendritic growth in sONα and tONα ganglion cells. **(A)** Schematic representation illustrating how input parameters (branching factor, growth radius, noise factor, and boundary tightness) influence dendritic morphology within the computational model. **(B)** Example model-generated dendritic reconstructions for sONα (top) and tONα (bottom) ganglion cells at different developmental stages, using optimized parameters that best fit biological growth patterns. Scale = 50 µm. **(C)** t-SNE visualization of model-generated (top) and real (bottom) dendritic reconstructions, showing clustering based on structural similarity. **(D)** Computational timelines mapping simulated developmental iterations to real postnatal ages. Size-related parameters are shown for sONα (left) and tONα (right), with the y-axis representing model iterations and the x-axis representing real developmental time points. **(E)** Sholl analysis comparing dendritic density distributions in real (blue) and model-generated (gray) ganglion cells. Example dendritic traces are shown for each developmental stage, illustrating morphological correspondence between real (blue, top) and simulated neurons (gray, bottom) at equivalent time points. **(F)** Cumulative plots illustrating the growth trajectories of different dendritic parameters in real (blue gradient) and model-generated (gray gradient) neurons. Lighter shades represent earlier developmental stages, while darker shades indicate later stages. sONα cells are displayed in the top plots, and tONα cells in the bottom plots.

Bayesian optimization refined four key parameters governing dendritic growth: the balancing factor (bf), which balances wiring length and path efficiency; the growth radius (r), which sets the maximum branch extension per step; the noise factor (k), which controls variability in branch placement; and boundary tightness (α), which determines how completely the dendritic tree fills the available space (Supplementary Figure 11A). With this parameterization, the model replicated the distinct growth patterns of sONα and tONα cells (Figure 7B). Despite differences in final morphology, both cell types followed the same underlying growth rules, as a single parameter set applied across all developmental stages (Supplementary Figure 11B).

Applying the optimized parameters across all developmental stages, we reconstructed synthetic dendritic trees. t-SNE analysis confirmed that synthetic neurons grouped closely with real ones, demonstrating that the model accurately captured biological dendritic structures (Figure 7C; Supplementary Figure 11 C-E). By reconstructing dendritic trees branch by branch, the model produced growth curves for the size parameters, establishing a computational timeline for dendritic development (Figure 7D).

Size-based parameters, e.g. total dendritic length and growth radius, followed a continuous trajectory, aligning well with biological development. In contrast, branching-based parameters, including branch points and dendritic intersections, remained stable across developmental stages, suggesting that branching is established early and maintained over time. Although the model followed the same growth rules for both cell types, the timing of development differed. sONα cells expanded more gradually than tONα cells, which exhibited rapid early-stage growth (Figure 7D). At P03, the iteration interval for sONα cells was 46.36 ± 25.86, while for tONα cells, it was 167.58 ± 34.14, reflecting faster initial expansion in tONα cells. This trend persisted, with tONα cells reaching higher iteration intervals across all developmental stages, reinforcing that sONα cells undergo prolonged growth, whereas tONα cells stabilize earlier.

The model’s predictions were further validated by comparing Sholl analyses between synthetic and biological dendritic trees (Figure 7E). Additionally, cumulative growth trajectories closely aligned, demonstrating the model’s ability to replicate cell-type-specific dendritic development (Figure 7F). This computational framework provides a quantitative approach for estimating a neuron’s developmental stage and offers a foundation for simulating dendritic responses to injury.

### Neuroprotective treatment induces further reduction in dendritic maturity

To investigate how promoting axon regeneration affects dendritic morphology, we activated the mTOR and JAK-STAT3 pathways by co-deleting the phosphatase and tensin homolog (PTEN) and the suppressor of cytokine signaling 3 (SOCS3) inhibitors in ganglion cells. This combined activation is well-known for promoting robust axon regeneration compared to the activation of each pathway independently (8,34,35). For this we used a dual-viral labeling strategy in PTEN^fl/fl^;SOCS3^fl/fl^ mice, that enabled sparse labeling of targeted ganglion cells while deleting PTEN and SOCS3. A retrograde Cre virus was injected into the superior colliculus (a major retinal target receiving input from ~85% of all ganglion cells), 14 days before injury to initiate recombination in retinal ganglion cells, followed by an intravitreal injection of a Cre-dependent AAV expressing tdTomato to visualize dendritic architecture (Figure 8A; see Methods). The dendritic structure was analyzed at 2 days post-injury (2 dpi) in three groups: uninjured controls (ctrl), optic nerve crush without regenerative treatment (injury), and ONC with conditional deletion of PTEN and SOCS3 (cdKO) (Figure 8B). This early time point was chosen to capture initial post-injury remodeling while regeneration was still ongoing (7).

**Figure 8.**
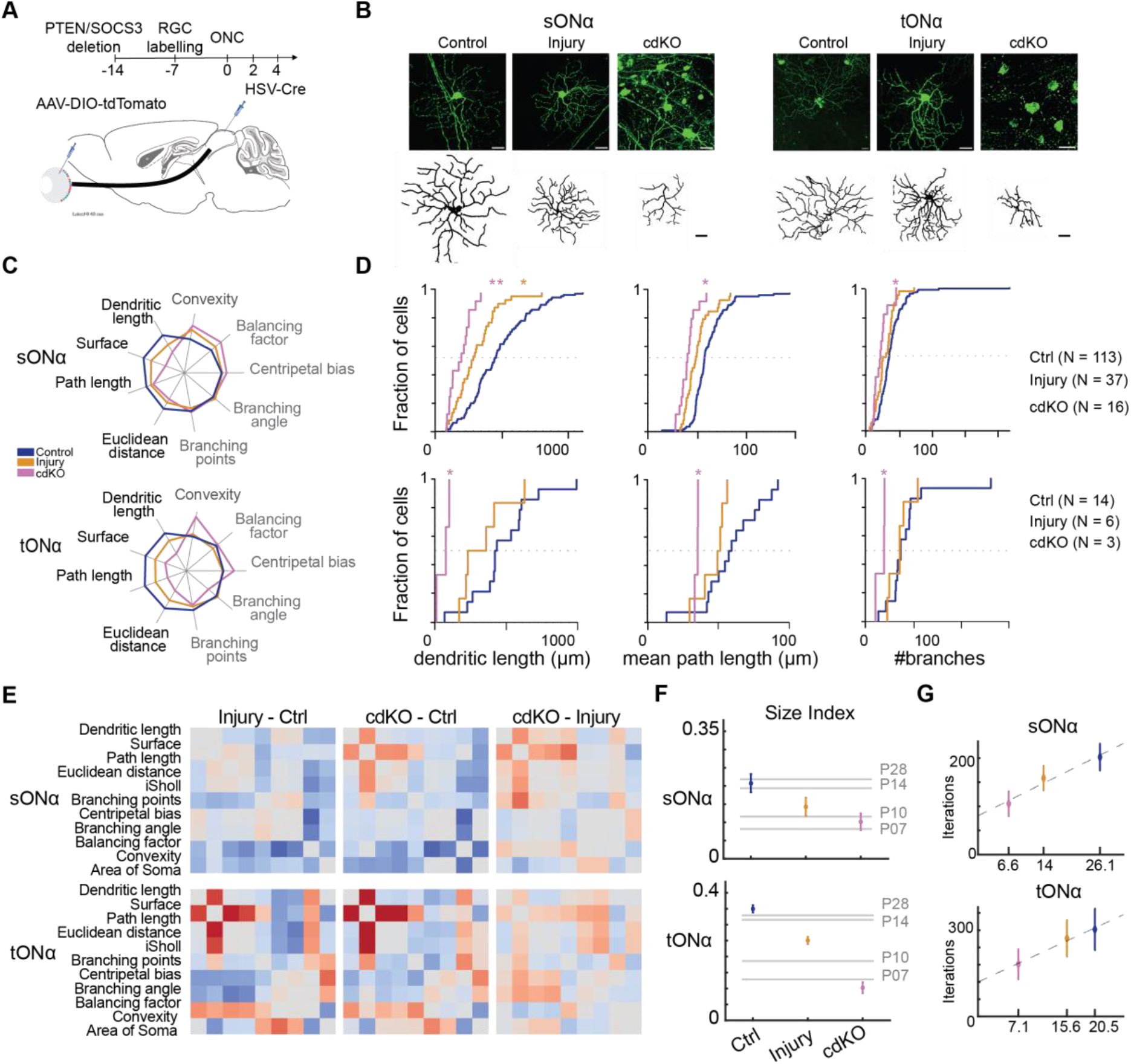
PTEN/SOCS3 treatment promotes additional shrinkage after injury. **(A)** Schematic representation of the experimental design. PTEN/SOCS3 were co-deleted in ganglion cells using a dual-viral strategy. The timeline indicates labeling, injury induction, and post-injury analysis. **(B)** Representative images of sONα (left) and tONα (right) ganglion cells across conditions: control (Ctrl), injury-only, and PTEN/SOCS3 co-deletion (cdKO). Top: En-face confocal images. Bottom: Corresponding traced dendritic reconstructions. Scale bars: 50 µm. **(C)** Quantification of dendritic morphology across conditions (Ctrl, Injury, cdKO). Left: size-related parameters (black). Right: branching metrics (light gray). **(D)** Cumulative distribution plots of total dendritic length, mean path length and number of branches for sONα (top row) and tONα (bottom row) across conditions. Statistical comparisons performed using KS test. **(E)** Differential Pearson correlation heatmaps showing structural shifts between experimental conditions. Each matrix represents the subtraction of a Pearson correlation matrix between two conditions: Injury – Ctrl (left), cdKO – Ctrl (middle), and cdKO – Injury (right). **(F)** Quantification of dendritic size across conditions. Colored error bars represent the mean and standard error of the mean (SEM) for each condition (Ctrl, Injury, cdKO). Horizontal gray lines indicate mean values for corresponding developmental stages. Top: sONα cells; Bottom: tONα cells. **(G)** Iteration-based timelines fitting of size parameters of experimental conditions to the computational model for sONα (top) and tONα (bottom). Each condition was aligned to the model to determine which iteration corresponded to the observed morphology. This allowed the estimation of a developmental equivalent timepoint for each condition.

PTEN/SOCS3 co-deletion exacerbates dendritic shrinkage after injury, further reducing dendritic size beyond what is observed in optic nerve crush alone. At 2 dpi, both sONα and tONα ganglion cells exhibited significantly reduced dendritic length compared to injured cells and uninjured controls (Figure 8C). In cdKO cells, dendritic length was 1663.99 ± 728.22 μm, compared to 2915.00 ± 1547.62 μm in injury and 4491.10 ± 2384.45 μm in uninjured controls. This effect was even more pronounced in tONα cells, where dendritic length was reduced to 1227.56 ± 387 μm in cdKO, compared to 3521.86 ± 1425.04 μm in injury and 4827.87 ± 1973.22 μm in uninjured controls (Figure 8D). Despite this shrinkage, dendritic branching features remained largely unchanged in sONα cells, with only minor variations in convexity and balancing factors. In contrast, tONα cells exhibited greater reductions in branching following treatment, suggesting structural reorganization beyond simple shrinkage.

Correlation analyses revealed distinct effects of PTEN/SOCS3 deletion on dendritic structure (Figure 8E). In sONα cells, dendritic morphology remained highly correlated with controls, despite size reduction (r = 0.882 for injury vs. control; r = 0.842 for cdKO vs. control). In contrast, tONα cells showed greater deviations from baseline, with correlation values dropping to r = 0.572 for injury vs. control and r = 0.563 for cdKO vs. control, indicating significant structural reorganization.

Parameter indices provided further insight into these differences (Figure 8F). These were calculated by normalizing parameters between 0 and 1 and grouping them by size-related features (e.g., dendritic length, mean path length) and branching-related features (e.g., branch order, density). In sONα cells, dendritic size decreased by 51.16% compared to uninjured controls (p = 0.014) and by 28.99% relative to injury (p = 0.242), reaching values typical of earlier postnatal stages (P10). tONα cells exhibited a more pronounced reduction, with a 59.17% decrease compared to injury and a 70.74% decrease relative to uninjured controls (p < 0.0001). Despite this reduction in dendritic size, branching remained largely stable in sONα cells, with no significant differences across conditions (p = 0.811 for cdKO vs. injury). In tONα cells, branching also remained mostly unchanged, though small differences in the cdKO group suggested slight remodeling rather than a complete loss of structural integrity.

Finally, computational modeling revealed that PTEN/SOCS3 deletion shifted both sONα and tONα cells toward earlier developmental morphologies (Figure 8G). In sONα cells, cdKO neurons aligned with P7.93 ± 2.98 (SEM), whereas injured cells corresponded to a later developmental stage, P12.42, interpolating between P10 and P14. Similarly, in tONα cells, injured neurons aligned with P12.83 ± 2.90 (SEM), but PTEN/SOCS3 deletion prevented further maturation, locking them in an earlier state. While both cell types regressed to similar developmental ages, prior modeling (Figure 7) suggests that this age corresponds to different stages of dendritic development in each cell type. In sONα cells, P10 represents an ongoing growth phase, while in tONα cells, it reflects a structurally stable state. This distinction may have implications for how each cell type responds to injury and regeneration.

## Discussion

Neuronal regeneration in the adult mammalian central nervous system is highly restricted, yet neurons retain developmental programs that could be reactivated to support repair. This study demonstrates that alpha ganglion cell resilience and regenerative potential are closely linked to their developmental growth trajectories. Alpha ganglion cells exhibit distinct maturation timelines, with tONα cells stabilizing by P10 and sONα cells continuing to grow until P14. These differences in developmental timing appear to shape their responses to both injury and regeneration, reinforcing the idea that intrinsic maturation dynamics influence long-term structural resilience.

Differences in the developmental maturation of sONα and tONα cells strongly influence their responses to optic nerve injury, with distinct patterns of dendritic regression emerging post-injury. The distinct maturation trajectories of sONα and tONα cells set the stage for their divergent responses to optic nerve injury.

Following optic nerve crush, sONα cells undergo rapid and structured dendritic shrinkage, whereas tONα cells exhibit delayed but more extensive regression. Despite these differences in timing, dendritic regression in both subtypes mirrors early developmental stages. sONα cells revert to P10-P14 morphologies through a controlled and structured process, while tONα cells display greater variability in their regression, reflecting a less coordinated adaptation. This variability may impact functional recovery, as unstructured shrinkage could compromise connectivity and signal integration, reinforcing the idea that developmental timing influences regenerative capacity. PTEN/SOCS3 co-deletion further intensifies these patterns, shifting both cell types toward even earlier developmental states (P07-P10). Notably, although both subtypes regress to similar modeled developmental ages, the functional implications differ: in sONα cells, P10 represents an ongoing phase of structural refinement, whereas in tONα cells, the same stage corresponds to a near-mature state. This difference underscores the importance of developmental timing in shaping injury responses and regenerative capacity.

These findings align with previous work suggesting that dendritic and axonal growth are highly interdependent during neuronal maturation (36). The earlier stabilization of tONα cells suggests a more constrained plasticity, potentially limiting their ability to respond dynamically to injury. Conversely, the prolonged growth phase of sONα cells may confer greater adaptability, allowing for more structured dendritic remodeling post-injury. Studies in neurodegeneration models further support these interpretations: sONα cells shrink rapidly but maintain dendritic organization, whereas tONα cells experience progressive and disorganized regression, a feature associated with greater vulnerability (6,37). Electrophysiological data indicate that despite substantial dendritic shrinkage, alpha ganglion cells retain core functional properties such as ON-OFF responses and orientation selectivity. This suggests that synaptic integrity is largely preserved even as dendritic architecture is lost. However, firing rates alone may not fully capture functional preservation. Future work employing calcium imaging, optogenetics, and synaptic connectivity analysis will be necessary to determine whether structural regression is truly decoupled from synaptic function or if compensatory mechanisms sustain network stability despite dendritic loss.

The interplay between dendritic remodeling and axonal regeneration remains a key question in neuronal recovery. PTEN/SOCS3 co-deletion enhances axonal regrowth but also exacerbates dendritic shrinkage. This dual effect suggests that dendritic regression may serve as an adaptive mechanism, reallocating resources toward axonal repair. Similar mechanisms have been described in zebrafish, where dendritic shrinkage occurs immediately after injury and appears to facilitate axonal regeneration rather than merely reflecting damage (15). In developing neurons, dendritic remodeling precedes the establishment of mature circuits, mirroring the post-injury regression observed here (38). The fact that inhibiting dendritic retraction in zebrafish delays axonal regrowth further supports the idea that temporary structural regression may be necessary to support axonal regeneration.

Building on these observations, our findings suggest that the effects of PTEN/SOCS3 co-deletion are cell-type dependent, further reinforcing the interplay between developmental timing and regenerative capacity. sONα cells, which regenerate more effectively, undergo controlled and organized dendritic shrinkage, preserving structural integrity even as their morphology shifts to an earlier state. In contrast, tONα cells, which exhibit a less coordinated response, appear to be structurally fixed in an arrested form, limiting their ability to further adapt. This suggests that sONα cells retain greater plasticity despite their regression, whereas tONα cells, due to their earlier developmental stabilization, may experience constrained remodeling. These distinctions reinforce the importance of developmental timing in shaping regenerative potential and highlight the need to consider intrinsic growth trajectories when designing targeted interventions. Recent work has shown that PTEN/SOCS3 deletion enhances axonal regeneration by increasing mitochondrial transport and glycolysis at regenerating axon tips (7). These metabolic shifts may explain why enhanced regeneration is accompanied by dendritic shrinkage: an energy trade-off that prioritizes axonal repair over dendritic maintenance.

These findings indicate that PTEN/SOCS3 deletion does not simply rewind development but resets each cell type to a distinct functional state, aligning with and expanding upon current theories of regenerative neurobiology. While many studies suggest that neuronal injury responses may involve the reactivation of developmental programs (39) our findings highlight that these programs do not operate uniformly across cell types. The observed differences in dendritic regression and structural reorganization suggest that regeneration is not merely a reversal of maturation but a nuanced process dependent on intrinsic developmental timing and plasticity. While sONα cells may preserve plasticity despite their regression, tONα cells appear to become structurally fixed in an incomplete form, limiting their ability to further adapt. The developmental timeline thus emerges as a key determinant of neuronal resilience: neurons that sustain prolonged growth retain greater regenerative potential, while those that stabilize early face greater constraints. Understanding these intrinsic differences will be essential for designing regenerative strategies that balance axonal outgrowth with dendritic preservation, ensuring both structural integrity and functional recovery.

Furthermore, this study provides novel insights into the dual roles of dendrites in neuronal injury and regeneration, emphasizing their active participation in neuronal recovery. Our findings suggest that dendrites are not merely passive victims of injury but may represent an adaptive response that reallocates cellular resources and activates developmental programs to support axonal regeneration.

## Acknowledgments

We thank Frederique Ooms for the animal maintenance and the technical support and Norma Kuhn for the critical discussions. This work was supported by the FWO (G091719N to KF, 12S7917N/12S7920N to KR, 1S42720N to LuM); the KU Leuven Research Council (C14/18/053 and C14_22_074 to KF and LM); and the NIH (1RO1EY032101 to KF).

## Author contributions

JRF and KF conceived the study. JRF, CL, LM, and KF designed the experiments. JRF performed surgeries/dissections, cell tracings, and modeling experiments. CL conducted the Santos et al., Sep 2025 – preprint copy – BioRxiv electrophysiology and two-photon calcium imaging experiments. LA performed the optic nerve crush experiment in PTEN/SOCS3 mice. JRF, CL, HC and KF analyzed and curated the data. JRF, KR, KF and HC developed the methodology. JRF, BN, KR, KF and HC developed the software. JRF, KF, and LM managed the project. Supervision was provided by LM, KF, and HC. JRF, CL, and KF prepared the visualizations. JRF, CL, and KF wrote the manuscript with input from LM, LuM, KR and HC

## Materials and Methods

**Resources Table**

| REAGENT OR RESOURCE | SOURCE | IDENTIFIER |
| --- | --- | --- |
| **Antibodies** |  |  |
| Anti-GFP (rabbit polyclonal) | Thermo Fisher Scientific | Cat# A-11122; RRID:AB_221569 |
| Anti-GFP (chicken polyclonal) | Thermo Fisher Scientific | Cat# A-10262; RRID:AB_2534023 |
| Anti-ChAT (goat polyclonal) | Millipore | Cat# AB144P; RRID:AB_2079751 |
| SMI32 (mouse monoclonal) | BioLegend | Cat# 801701; RRID:AB_2564642 |
| Anti-CART (rabbit polyclonal) | Phoenix Pharmaceuticals | Cat# H-003-62; RRID:AB_2313614 |
| Anti-FOXP2 (goat polyclonal) | Abcam | Cat# ab1307; RRID:AB_1268914 |
| Anti-mCherry (donkey anti-rabbit) | Novus Biologicals | Cat# NBP2-25158; RRID:AB_2636881 |
| Alexa Fluor 488 donkey anti-rabbit | Thermo Fisher Scientific | Cat# A-21206; RRID:AB_2535792 |
| Alexa Fluor 488 donkey anti-chicken | Jackson ImmunoResearch | Cat# 703-545-155; RRID:AB_2340375 |
| Alexa Fluor 633 donkey anti-goat | Thermo Fisher Scientific | Cat# A-21082; RRID:AB_10562400 |
| Cy3 donkey anti-mouse | Jackson ImmunoResearch | Cat# 715-165-151; RRID:AB_2315777 |
| Alexa Fluor 555 donkey anti-goat | Abcam | Cat# ab150130 |
| DyLight 405 donkey anti-rabbit | Jackson ImmunoResearch | Cat# 711-475-152; RRID:AB_2340839 |
| Cy3 donkey anti-chicken | Jackson ImmunoResearch | Cat# 703-166-155; RRID:AB_2340364 |
| **Virus strains** |  |  |
| HSV-Cre (Herpes Simplex Virus type 1 expressing Cre recombinase under hEF1 promoter) | Massachusetts Institute of Technology | RN431 hEF1-Cre HT |
| HSV-Flp (Herpes Simplex Virus type 1 expressing Flpo recombinase under hEF1 promoter.) | Massachusetts Institute of Technology | RN433 hEF1-Flpo HT |
| AAV2.2-CAG-DIO-mitoGFP-t2A-tdTomato-WPRE | Custom construct | Not specified |
| AAV2.2-CAG-fDIO-mitoGFP-t2A-tdTomato-WPRE | Custom construct | Not specified |
| **Organisms/strains (*Mus musculus*)** |  |  |
| Thy1-YFP-H | The Jackson Laboratory | JAX: 003782 |
| PTEN<sup>fl/fl</sup> | The Jackson Laboratory | JAX: 004597 |
| SOCS3<sup>fl/fl</sup> | The Jackson Laboratory | JAX: 010944 |
| **Chemicals, peptides, and recombinant proteins** |  |  |
| 10x PBS | VWR | Cat# 437117K |
| 1x PBS | VWR | Cat# 444057Y |
| Histofix 4% | Roche | Cat# P087.5 |
| Normal Donkey Serum | Millipore | Cat# S30-100ML |
| 10% Bovine Albumin | Sigma | Cat# SRE0036-250ML |
| DMEM, high-glucose | Thermo Fisher Scientific | Cat# 41965062 |
| Trypsin 0.05% | Thermo Fisher Scientific | Cat# 25300054 |
| Fetal Bovine Serum (FBS) | Thermo Fisher Scientific | Cat# 10270106 |
| 2,2'-thiodiethanol (TDE) | Sigma | Cat# 166782-500G |
| ProLong Gold Antifade Mounting Medium | Thermo Fisher Scientific | Cat# P36934 |
| Sodium Azide (NaN<sub>3</sub>) | Sigma | Cat# S2002-100G |
| Triton X-100 | Sigma | Cat# T8787 |
| Sodium Chloride (NaCl) | Sigma | Cat# S7653-250G |
| Potassium Chloride (KCl) | Sigma | Cat# P5405-25G |
| Calcium Chloride (CaCl<sub>2</sub>) | Sigma | Cat# C5670-100G |
| Magnesium Chloride (MgCl<sub>2</sub>) | Sigma | Cat# M8266 |
| D-glucose (Dextrose) | Sigma | Cat# G8270-250G |
| Sodium phosphate monobasic (NaH<sub>2</sub>PO<sub>4</sub>) | Sigma | Cat# S5011 |
| Sodium hydroxide (NaOH) | Sigma | Cat# 655104-500G |
| Sodium bicarbonate (NaHCO<sub>3</sub>) | Sigma | Cat# S8875-1KG |
| **Software and algorithms** |  |  |
| Fiji | https://fiji.sc/ | RRID:SCR_002285 |
| MATLAB | MathWorks | RRID:SCR_001622 |
| ZEN Lite | Zeiss |  |
| ChAT band detector | This paper | https://github.com/farrowlab/ChATbandsDetection_DLC |
| Python 3.4 | Python Software Foundation | www.python.org |
| TREES toolbox | Cuntz et al. (2011) | https://github.com/cuntzlab/ |
| Psychophysics Toolbox | Psychtoolbox | http://psychtoolbox.org/ |
| WaveSurfer (version: 0.918) | Janelia Research Campus | http://wavesurfer.janelia.org/ |
| ScanImage | Former: Vidrio Technologies, now: MBF Bioscience | https://docs.scanimage.org/ |
| **Other** |  |  |
| Wiretrol II capillary micropipettes | Drummond Scientific | Cat# 5-000-2005 |
| Borosilicate glass | Sutter Instrument | Cat# BF100-20-10 |
| Rapid Flow Filters 0.2 µm pore size | VWR | Cat#. 514-0027 |
| Laser-based Micropipette Puller | Sutter Instrument | Cat# P-2000 |
| Rapid Flow Filters 0.2 mm pore size | VWR | Cat# 514-0027 |
| Premium Standard Wall Borosilicate capillary glass | Warner Instrument | Cat# G100-4 |
| Small Animal Stereotaxic Workstation | Narishige | Cat# SR-5N |
| Stereotaxic Micromanipulator | Narishige | Cat# SM-15R |
| Hydraulic Oil Micromanipulator | Narishige | Cat# MO-10 |
| Oil Microinjector | Narishige | Cat# IM-8B |
| Two-photon microscope | Scientifica | Serial# 14200 |
| 780 nm LED light source | Thorlabs | Cat# M780L3 |
| Patch-Clamp amplifier | Molecular Devices | Axon Multiclamp 700B |
| Patch-Clamp microscope | Scientifica | Slice Scope |
| Patch-Clamp manipulator | Scientifica | Serial# 301311 |
| Zeiss LSM 710 confocal microscope | Zeiss | Cat# LSM710 |

### Mice

All experimental procedures were approved by the Ethical Committee for Animal Experimentation (ECD) of the KU Leuven and followed the European Communities Guidelines on the Care and Use of Laboratory Animals (166-2018/EEC, 168-2022/EEC). Both male and female adult (0-3 months old) Thy1-YFP-H mice and PTEN^fl/fl^SOCS3^fl/fl^ were used in our experiments. Mice were kept on a 12-h light-dark cycle (lights on at 7:00), and sterilized food pellets and water were provided *ad libitum*.

### Tissue harvest (retina)

Mice were sacrificed at postnatal days P03, P07, P10, P14, P28, and at 0, 1, 2, 4, 7, and 14 days post-injury (6–8 weeks old) following protocols approved by the institutional animal care and use committee. Deep anesthesia was induced using an intraperitoneal injection of Ketamine/Xylazine/Acepromazine (100/10/3 mg/kg). Retinas were carefully dissected and post-fixed in 4% PFA at room temperature for 2 hours. Post fixation, tissues were rinsed thoroughly in PBS, and retinas were separated for further processing. This method ensured optimal preservation of tissue morphology for downstream analyses.

### Surgical procedures

Animals were anesthetized using Isoflurane (Iso-vet 1000 mg/mL) to achieve rapid sedation, followed by an intraperitoneal injection of Ketamine and Medetomidine (0.75 mL Ketamine, 100 mg/mL; 1 mL Medetomidine, 1 mg/mL; diluted in 8.2 mL saline). Mice were placed in a stereotaxic workstation (Narishige, SR-5N) to ensure precise surgical positioning. To protect the eyes during the procedure, Dura-tear (NOVARTIS, 288/28062–7) was applied.

Surgical labeling of retinal ganglion cells was conducted on PTEN^fl^/SOCS3^fl^ mice. A Cre-dependent herpes-simplex virus (HSV) was injected into the superior colliculus to co-delete regenerative proteins in the regeneration group, while a FLP-dependent HSV was used for the control and injured groups. HSV injections were performed using Wiretrol II capillary micropipettes (Drummond Scientific, 5-000-2005) with an open tip (~30 µm) and an oil-based hydraulic micromanipulator (Narishige MO-10).

Injection coordinates for 4-week-old mice (bregma-lambda distance of 4.7 mm) were adjusted based on individual anatomical variations. For a typical mouse, stereotactic coordinates were set as follows: anterior-posterior (AP): −4.20 mm, mediolateral (ML): ±1.95 mm, dorsoventral (DV): 3.50 mm. A total of 100–400 nL HSV was injected in single doses (up to 200 nL), with a waiting period of 5–10 minutes between each injection to minimize tissue displacement.

Fourteen days post-HSV injection, an adeno-associated virus (AAV) was delivered intravitreally to label ganglion cells. To maximize the randomization of ganglion cell labeling, HSV was injected into the superficial layers of the superior colliculus at a depth of 1.7–1.8 mm across four locations within a 1 mm^2^ field anterior to lambda, starting at the midline. Each injection delivered 100–200 nL of HSV, ensuring comprehensive coverage of the targeted area.

### Optic nerve crush

Optic nerve crush (ONC) was performed on mice (N = 5–7 per timepoint) (40). Mice were anesthetized via intraperitoneal injection of Ketamine (100 mg/kg) and Xylazine (10 mg/kg). The right eye of each mouse was used for the ONC procedure. The conjunctiva was carefully incised at the lateral canthus region and gently peeled back to expose the optic nerve. A small surgical window was created between the surrounding muscles to access the optic nerve approximately 0.5 mm posterior to the globe. The nerve was crushed for 4 seconds using self-closing jeweler’s forceps, ensuring consistent application of pressure while avoiding damage to surrounding muscles or blood vessels to prevent retinal ischemia. injured animals of the same strains served as controls for each experimental group.

### Retina immunohistochemistry

Mouse retinas were processed using immunohistochemistry protocols to assess dendritic morphology and markers such as SMI32 and CART. Dissected retinas, prepared as described previously, underwent fixation in 4% paraformaldehyde (PFA; Histofix, ROTH, P087.5 mm) supplemented with 100 mM sucrose for 30 minutes at 4°C. Following fixation, retinas were washed three times in 1x PBS (10 minutes each) at room temperature, cryoprotected by sequential incubation in 10%, 20%, and 30% sucrose in 1x PBS containing 0.1% sodium azide (NaN_3_), and stored overnight at 4°C. Freeze-cracking was performed by freezing retinas on slides coated with 30% sucrose for 3–5 minutes on dry ice, followed by thawing at room temperature. This freeze–thaw cycle was repeated twice to enhance tissue permeability. Retinas were washed three times in 1× PBS, then incubated in blocking buffer (10% normal donkey serum (NDS), 1% bovine serum albumin (BSA), 0.5% Triton X-100, and 0.02% NaN_3_ in 1× PBS) for at least 1 hour at room temperature.

Primary antibody staining was performed over 5–7 days at room temperature with gentle shaking. The primary antibody solutions included combinations of rabbit anti-GFP (Invitrogen, A-11122, 1:500), goat anti-ChAT (Chemicon, Ab144P, 1:200), chicken anti-GFP (Invitrogen, A-10262, 1:500), mouse SMI32 (Biolend, 801701, 1:1000), and rabbit anti-CART (Phoenix, H-003–62, 1:500), prepared in 3% NDS, 1% BSA, 0.5% Triton X-100, and 0.02% NaN_3_ in 1× PBS.

Following primary incubation, retinas were washed three times (10–15 minutes each) in 1× PBS with 0.5% Triton X-100. Secondary antibody staining was then performed using Alexa488 donkey anti-rabbit (Invitrogen, A-21206, 1:500), Alexa488 donkey anti-chicken (ImmunoJackson, 703-545-155, 1:500), Alexa633 donkey anti-goat (Invitrogen, A-21082, 1:500), Cy3 donkey anti-mouse (ImmunoJackson, 715-165-151, 1:400), and DyLight 405 donkey anti-rabbit (ImmunoJackson, 715-475-150, 1:200), prepared in 3% NDS, 1% BSA, 0.5% Triton X-100, and 0.02% NaN_3_ in 1× PBS. Nuclei were stained simultaneously with DAPI (Roche, 10236276001, 1:500). Retinas were incubated in secondary antibody solutions overnight at 4°C.

Following staining, retinas were washed three times in 1× PBS with 0.5% Triton X-100 and once in 1× PBS. Mounting was Santos et al., Sep 2025 – preprint copy – BioRxiv performed by sequential immersion of retinas in 10%, 25%, 50%, and 97% 2,2′-Thiodiethanol (TDE; Sigma, 166782–500G; (41)) for at least 30 minutes each, followed by embedding in ProLong Gold Antifade Mountant (Thermo, P36934) under #0 coverslips (MARIENFEL, 0100032, No.0, 18 × 18 mm). To prevent compression, four strips of Parafilm (PM999) were placed around the retinas before applying the coverslip. Alternatively, some retinas were mounted in 97% TDE with DABCO (Sigma, 290734) after immersion in TDE. The coverslips were sealed with nail polish to prevent evaporation, and samples were stored in the dark at 4°C.

### Confocal microscopy

Confocal microscopy was performed using a Zeiss LSM 710 microscope to capture high-resolution images of ganglion cellsand overall retinal structure. Overview images of the retina and brain were acquired using a 10× plan-APOCHROMAT objective (numerical aperture (NA): 0.45; Zeiss). Imaging was conducted with a zoom factor of 0.7, utilizing a 5 × 5 tile arrangement with 0–15% overlap between tiles, achieving a resolution of 2.37 µm/pixel.

For detailed scanning of individual ganglion cells, a 63× plan-APOCHROMAT objective (NA: 1.4; Zeiss) was used. Scans were performed at a zoom factor of 0.7 with a 2 × 2 tile arrangement (tile overlap: 15%) to ensure complete coverage of cell structures. This setup provided an XY-resolution of 0.38 µm/pixel and a Z-resolution between 0.25 and 0.35 µm/pixel. Z-stacks captured depth profiles of up to 50 µm, allowing comprehensive three-dimensional reconstruction of the dendritic morphology. These imaging parameters ensured accurate visualization and quantification of both retinal ganglion cell structure and broader retinal features.

### Classification and stratification of retina ganglion cell subtypes

The dendritic morphology of GFP-stained retinal ganglion cells was determined following the same procedure as in ***(42)***. Briefly, confocal z-stacks of individual ganglion cells of anti-ChAT–and anti-GFP–stained retinas were denoised, down-sampled, and binarized to extract the position of the On- and Off-ChAT bands using a custom deep learning framework and the position of the dendritic tree (anti-GFP). The dendritic trees were traced with TREES Toolbox ***(25)***. Z-stacks were warped to straighten the ChAT bands and a dendritic depth profile was determined from the number of bright pixels of the binarized GFP signal in the Z-direction. Dendritic tree diameters were calculated from the area of the convex hull of the dendritic tree in the X-Y plane as D =2*(area/π)1/2.

The density of dendrites was plotted along the Z-axis. Each cell was classified as below, between, or above ChAT bands based on dendritic stratification depth. The bistratified cells were with dendritic density peaks aligned with the ChAT band. (Github: https://github.com/farrowlab/Reinhard_2019).

### Extraction of ChAT-positions

ON and OFF ChAT bands were automatically localized in retinal slices using a modified version of DeepLabCut. The model’s deconvolutional layer was adapted to predict band locations only within the image’s center column. The model was subsequently applied to 2D slices using a sliding window approach (20 µm width, 10 µm step size), predicting the location of the ON and OFF bands in the center column of each window. Training data consisted of ON and OFF bands that were manually labeled at an approximate spacing of 20 µm using Fiji.

(Github: https://github.com/farrowlab/ChATbandsDetection_DLC)

### Soma position and removal of noise

After warping, the soma position was determined by filtering the GFP-signal with a circular kernel (42). If this method detected the soma, it was used to remove the soma from the GFP-data and the center of mass was taken as the soma position. If this automated method failed, the soma position was marked manually. Afterwards, dendrites of other cells, axons, and noise were removed manually: The warped GFP-signal was plotted in side-view and en-face view in MATLAB, and pixels belonging to the cell were selected manually.

### Computation of the dendritic statistics

To compute the dendritic statistics a minimal spanning tree model was created of each imaged dendritic tree using the TREES toolbox with a balancing factor of 0.2. From this tree we calculated a set of five statistics including: the mean ratio of path length and Euclidean distance; maximum metric path length; mean branch lengths; mean path length and z-range against width of spanning field (25).

### Dendritic morphology tracing and parameter extraction

Dendritic reconstructions and parameter extractions were performed using the TREES Toolbox version 1.16 (25) for MATLAB. High-resolution confocal microscopy images were used to manually or semi-automatically trace dendritic arbors, which were subsequently resampled to a uniform internode distance of 1 µm to standardize the analyses.

### Neuronal Morphology Analysis

This study examines neuronal morphology using key quantitative parameters to characterize dendritic structures, spatial organization, and their functional implications. Understanding these metrics helps in analyzing neuronal connectivity, synaptic integration, and the structural branching of dendritic trees.

1. <u>Total Dendritic Length</u> represents the sum of all internode segment lengths in a neuron, capturing the full extent of its dendritic arbor.
2. <u>Number of Branch Points</u> quantifies the number of bifurcation points where a dendritic segment splits into two or more branches. It serves as an important indicator of dendritic branching features, influencing signal integration and transmission across the neuron.
3. <u>Surface Area</u>. The dendritic membrane’s total surface area and volume are essential for estimating synaptic density and integration potential. These parameters help in understanding the three-dimensional expansion of neuronal structures and their ability to interact with surrounding synaptic inputs.
4. <u>Soma Size</u>. The size of the soma is a fundamental characteristic that influences neuronal function. It is manually delineated from raw images.
5. <u>Sholl Analysis</u> evaluates the number of intersections between dendrites and a series of concentric circles centered on the soma.
  - <u>Integral Sholl Profile (iSholl):</u> The iSholl metric normalizes intersections to the integral of the Sholl profile, capturing dendritic density and spatial distribution (30)
6. <u>Path & Spatial Measures</u>
  - <u>Mean Path Length:</u> The average path distance from the soma to all terminal branches, reflecting how far electrical signals must travel.
  - <u>Mean Euclidean Distance</u>: Measures the straight-line distance between the soma and terminal dendritic points, providing a baseline estimate of dendritic reach.
  - <u>Theta:</u> The angular distribution of branches relative to a reference axis, aiding in the assessment of directional biases in dendritic growth.
  - <u>Radius:</u> The radial distance from the soma to a specific dendritic segment, often used in conjunction with Sholl analysis.
7. <u>Morphological Features</u>
  - <u>Balancing Factor:</u> This metric evaluates how evenly the dendritic structure is distributed, providing insight into neuronal organization and functional efficiency.
  - <u>Centripetal Bias:</u> Measures whether dendritic growth favors a specific direction, offering insight into growth constraints and network connectivity.
  - <u>Convexity:</u> Quantifies how much of the dendritic tree is enclosed within a convex hull. A higher convexity suggests a compact structure, whereas lower convexity indicates a more dispersed dendritic layout.
  - <u>Branching Angle:</u> The mean angle formed between dendritic bifurcations, influencing signal propagation and spatial organization.
8. <u>Additional Metrics</u>
  - <u>Branch Order Distributions:</u> Categorizes nodes and dendritic cable lengths based on their branch order, which increases at each bifurcation.
  - <u>Branch Length Distributions:</u> Measures path lengths from each node to the soma, dividing the tree into individual branches to assess variability in segment length.

These metrics were extracted from individual dendritic reconstructions to quantitatively describe the morphology of retinal ganglion cells across experimental conditions. All data were normalized to minimize inter-animal variability and ensure comparability.

### Computational Modeling of Dendritic Growth and Injury Responses

A computational model adapted from (33) was used to simulate dendritic growth and analyze injury-induced changes in retinal ganglion cells. The model employed a stretch-and-fill algorithm to generate synthetic dendritic trees that balanced wiring cost and spatial coverage. It was designed for Class IV fly larva dendritic arborization neurons but was shown to be a good match also for Class I and III neurons (43,44) as well as many other cell types including rodent cortex and hippocampus. Model parameters were optimized separately to sONα and tONα ganglion cells to match experimental data, according to the metrics mentioned above (total dendritic length, mean path length, mean branch order, convexity, theta, mean Euclidean distance, and branching angle).

The following key model parameters for the growth_tree function were fitted:

- <u>Balancing Factor (bf):</u> Optimized to 0.24 for sONα cells and 0.25 for tONα cells, controlling the trade-off between wiring cost and dendritic density.
- <u>Target Radius (r):</u> Set to 65.72 µm for sONα cells and 67.54 µm for tONα cells, defining the expected spatial extent of dendritic growth.
- <u>Branching Probability (k):</u> Adjusted to 0.65 for both sONα and tONα cells, governing the likelihood of new branch formation.
- <u>Asymmetry Parameter (a):</u> Adjusted to 0.66 for sONα cells and 0.67 for tONα cells, influencing the directional bias in dendritic branching.

### Bayesian Optimization Approach

Bayesian Optimization was used to identify the optimal parameters by minimizing the discrepancy between real and synthetic dendritic trees. The objective function compared generated dendritic trees against experimental data using:

- Principal Component Analysis (PCA): Reducing high-dimensional morphological metrics into key components for distance-based evaluation.
- Mahalanobis Distance: A statistical measure comparing the synthetic and real tree distributions.
- Cross-Correlation of Sholl Profiles: Evaluating the similarity of branching distributions.

This method ensures that the generated neuronal trees closely resemble experimentally observed structures, preserving dendritic architecture and connectivity patterns. These metrics and optimization strategies provide a standardized and detailed framework for analyzing dendritic morphology, ensuring consistency in assessing neuronal structure across different conditions and experimental models.

### Model Validation and Growth curves

Synthetic trees were validated against experimental reconstructions by comparing branching metrics using t-distributed stochastic neighbor embedding (t-SNE) and Sholl analyses, demonstrating high morphological alignment.

Developmental timelines generated by the model reflected progressive changes in size metrics (Total dendritic length, Surface, iSholl, # Branching points, Area of soma, Mean path length and mean Euclidean distance). Simulations of dendritic regression following injury showed morphological profiles resembling earlier developmental stages, aligning injured cells approximately with P14 profiles for both tONα and sONα cells. Additionally, cdKO simulations showed dendritic regression similar to earlier developmental stages, aligning injured cells approximately with P07 profiles. These findings highlight the utility of computational modeling for understanding injury-induced changes in Ganglion cells and predicting neuronal adaptation over time.

### Retinal electrophysiology

#### Preparation of retinas

Retinas were prepared from Thy1-YFP-H mice, which sparsely label retinal ganglion cells. Retina isolation was performed under deep red illumination to minimize photoreceptor activation. Ringer’s medium was used during dissection and recordings, containing 110 mM NaCl, 2.5 mM KCl, 1 mM CaCl2, 1.6 mM MgCl2, 10 mM D-glucose, and 22 mM NaHCO_3_, bubbled continuously with 5% CO2/95% O2 to maintain a physiological pH of 7.4.

The retinas were mounted with the ganglion cell-side up on filter paper (Millipore, HAWP01300) with a rectangular aperture (3.5 mm wide) in the center, ensuring proper exposure for electrophysiological recordings. The mounted retinas were then superfused with Ringer’s medium heated to 32–36°C in the microscope chamber during the experiments.

### Electrophysiology

Electrophysiological recordings were performed using an Axon Multiclamp 700B amplifier (Molecular Devices). Borosilicate glass electrodes (BF100-50-10, Sutter Instrument) were used with a resistance of 3–5 MΩ, filled with Ringer’s medium containing Alexa 555 for pipette visualization. Signals were digitized at 20 kHz using National Instruments hardware and acquired with WaveSurfer software (version 0.918) written in MATLAB. Recordings were conducted in loose cell-attached mode using the patch clamp technique to measure spiking activity. The setup was optimized for stability and minimal disruption to the neuronal environment. To visualize the pipette, Alexa 555 was added to the Ringer’s medium.

#### Targeted recordings using two-photon microscopy

Thy1-YFP positive cells were targeted for recording using a two-photon microscope (Scientifica) equipped with a Mai Tai HP two-photon laser (Spectra Physics) integrated into the electrophysiological setup. To facilitate targeting, two-photon fluorescent images were overlaid with the IR image acquired through a CCD camera. Infrared light was produced using the light from an LED. For some cells, z-stacks were acquired using ScanImage (Vidrio Technologies)

#### Presentation of visual stimuli

Visual stimuli were generated using a Samsung SP F10M LCD projector with a refresh rate of 60 Hz. The projector emitted light ranging from ~430 nm to ~670 nm, delivering a power density of 240 mW/cm^2^ at the retina. Stimulus intensity was adjusted using neutral density filters, reducing intensity by 1–2 log units. Custom software written in Octave, based on Psychtoolbox, controlled stimulus generation and presentation.

The following visual stimuli were presented to the retinas:

- <u>Full-field ‘chirp’ Modulation:</u> A stimulus with gray-black-gray-white-gray transitions (3 s per level), followed by a temporal modulation from black to white at increasing frequencies (0.5 Hz to 8 Hz over 6 s). After
- 3 s at gray, contrast modulation (0–100%) was performed over 5.5 s at 2 Hz. This sequence was repeated 10 times (32).
- <u>Spot-Size Stimuli:</u> Black or white spots of six different sizes (4°, 8°, 12°, 16°, 20°, 40°) were presented at the center of a gray screen for 2 s. Colors and sizes were randomized.
- <u>Large Moving Bar:</u> A black bar (40° wide) moved across the screen at 150°/s in eight directions (0°, 45°, 90°, 135°, 180°, 225°, 270°, 315°). Each direction was repeated five times in a randomized order.
- <u>Expansion Stimuli:</u> A black disc expanded linearly from 2° to 50° in diameter within 300 ms (150°/s) at the center of a gray screen. The stimulus was repeated 10 times.
- <u>White Expansion Stimuli:</u> A white disc expanded linearly from 2° to 50° in diameter within 300 ms (150°/s) at the center of a black screen. The stimulus was repeated 10 times.
- <u>Drifting gratings:</u> Drifting square-wave gratings with different spatial frequencies and contrast levels were shown in eight directions of random order in an aperture of 300 μm diameter on a dark background.

## Analysis of patch recordings

The loose-patch extracellular recording traces were high-pass filtered. Events that exceeded an amplitude threshold were extracted. Unless otherwise noted, firing rates were calculated as the number of spikes in 50 ms bins averaged across the 5–10 stimulus repetitions. Analysis of patch-clamp recordings was used the same procedure as in Reinhard et al. 2019. The repeats of visual stimuli and retinal responses were aligned using recorded stimulus start and frame triggers.

## Direction and orientation selectivity

of the ganglion cells was analyzed by calculating the DSI and OSI, defined as the vector sum of these peak responses for each of the eight different directions of the moving bar stimulus.

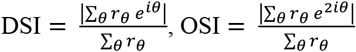

## On-off bias index

To estimate how the ganglion cell responds to increases or decreases in light intensity, we calculated an on-off bias index. This index compares the cell’s response to the spot of optimal size after the onset of ON and OFF flashes.

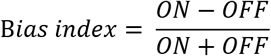

## Contrast tuning

To test the cells’ tuning to contrast, we used drifting gratings with optimal spatial tuning at different contrast levels: 25%, 50%, and 100%.

## Statistics

All statistical analyses were performed in MATLAB and Pyhton using built-in functions and custom scripts. The normality of the data was assessed using Shapiro-Wilk tests before choosing appropriate statistical methods. Comparisons between the two groups were conducted using the Mann-Whitney U test (Wilcoxon rank-sum test) for non-normally distributed data. The Kolmogorov-Smirnov (KS) test was used to compare distributions of morphological parameters across developmental and post-injury conditions. For correlation analyses, Pearson’s correlation was applied when data were linear and normally distributed; otherwise, Spearman’s rank correlation was used. The 2D Kolmogorov-Smirnov test was employed to assess differences in joint distributions of dendritic morphology parameters. To account for multiple comparisons, Benjamini-Yekutieli correction was applied to control the false discovery rate. For computational modeling, Bayesian optimization was used to estimate dendritic growth parameters, with prior distributions selected based on experimental data. Model convergence was assessed using posterior distributions and iteration stability. All tests were two-tailed, and significance was set at p < 0.05. Unless otherwise stated, data are presented as median ± interquartile range (IQR).

## Supplementary Figures

**Supplementary Figure 1.**
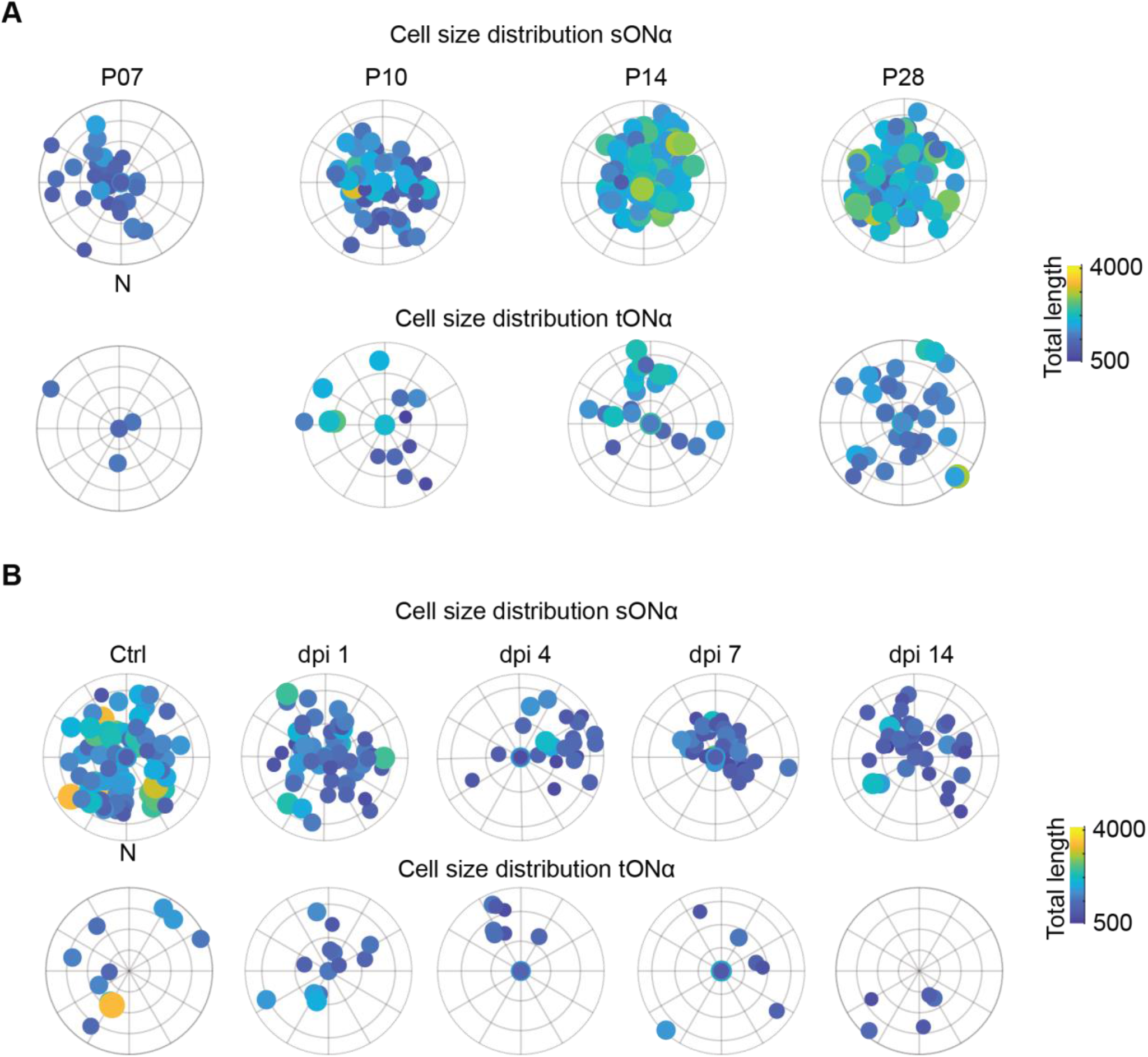
Polar plots of cell size distribution. **(A)** Cell size distribution during development. Polar scatter plots of sONα (top row) and tONα (bottom row) cells at different developmental stages (P07, P10, P14, P28). Similar to (A), dot size reflects cell area and color represents dendritic length, illustrating changes in spatial organization over time. The color scale represents the total dendritic length, with shorter trees in blue and longer trees in yellow. **(B)** Cell size distribution after injury. Polar scatter plots showing the spatial distribution of sONα (top row) and tONα (bottom row) retinal ganglion cells at different time points post-injury (Ctrl, dpi1, dpi4, dpi7, dpi14). Each dot represents a single cell, with dot size indicating cell area and color representing total dendritic length (see color bar). N denotes nasal orientation.

**Supplementary Figure 2.**
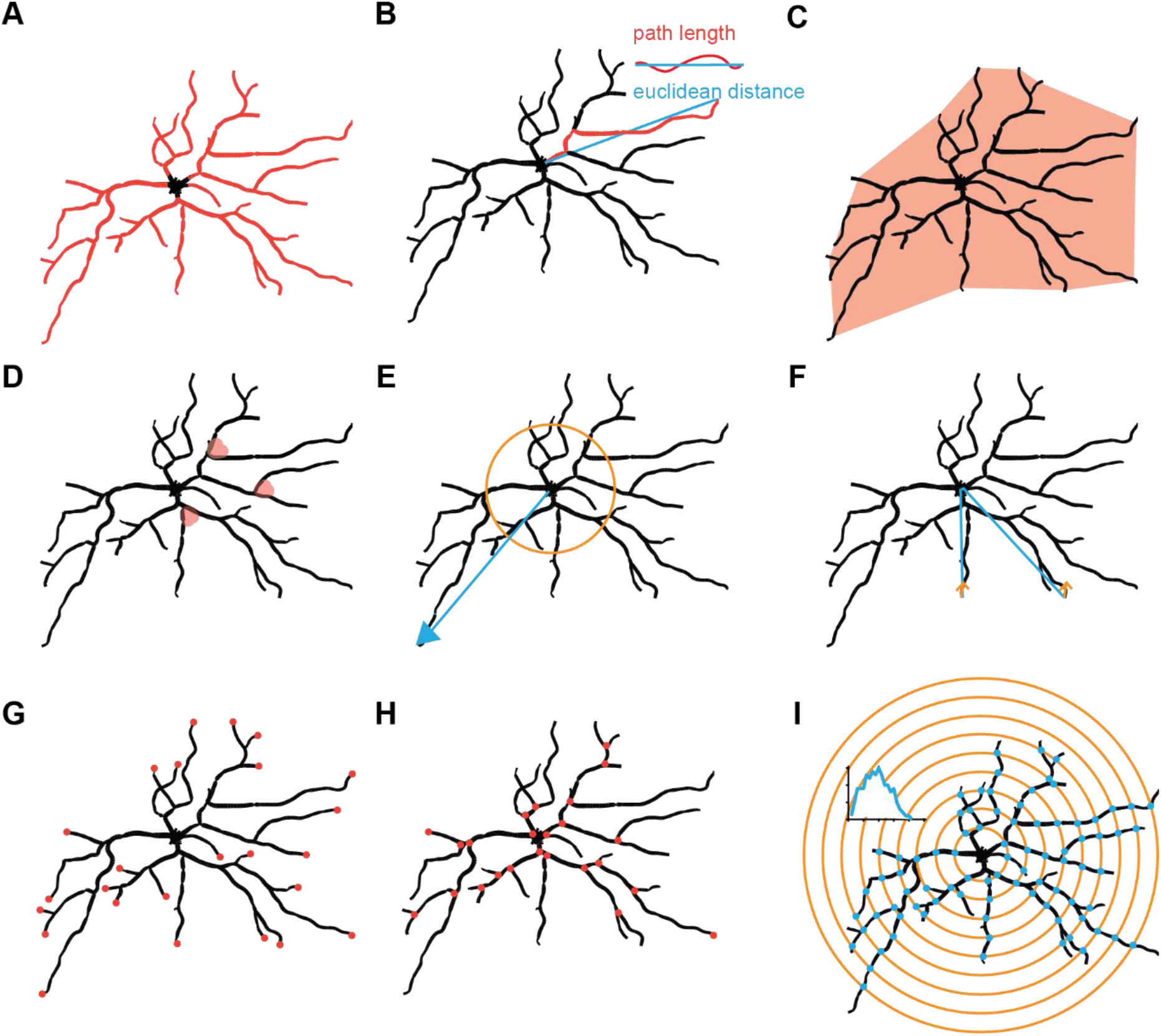
Representation of the dendritic morphological metrics. **(A)** Total dendritic length diagram. Total dendritic length is the sum of all dendritic segment lengths, representing the overall size of the arbor in microns. **(B)** Path length vs. Euclidean distance diagram. This plot compares the actual path length along dendritic branches (red) to the straight-line Euclidean distance (blue) between two points, highlighting the tortuosity of the dendritic tree. **(C)** Hull surface diagram. The convex hull (light coral) represents the smallest polygon enclosing all branch points, providing a measure of the spatial extent of the dendritic arbor. **(D)** Branching angle diagram. Branching angles represent the angular separation between two daughter branches at a bifurcation point, capturing structural complexity. **(E)** Root angle diagram. Root angles describe the initial directional bias of primary dendrites emerging from the soma, measured as the angle between the soma and the first bifurcation. **(F)** Centripetal bias diagram. Centripetal bias quantifies the directional tendency of dendritic growth, with blue arrows indicating the overall directional preference of branching toward or away from the soma. **(G)** Terminal branches diagram. Each red dot represents a terminal dendritic tip, marking the endpoints of dendritic paths and defining the outer extent of the arbor. **(H)** Branching points diagram. Each red dot represents a dendritic bifurcation, indicating locations where branches split and contributing to overall arbor complexity. **(I)** Sholl analysis diagram. Sholl analysis quantifies dendritic complexity by counting the number of branch intersections (teal dots) at evenly spaced distances from the soma (coral rings). The representative Sholl plot in the left corner shows the intersection count (teal) as a function of radial distance from the soma, illustrating how dendritic complexity changes with distance.

**Supplementary Figure 3.**
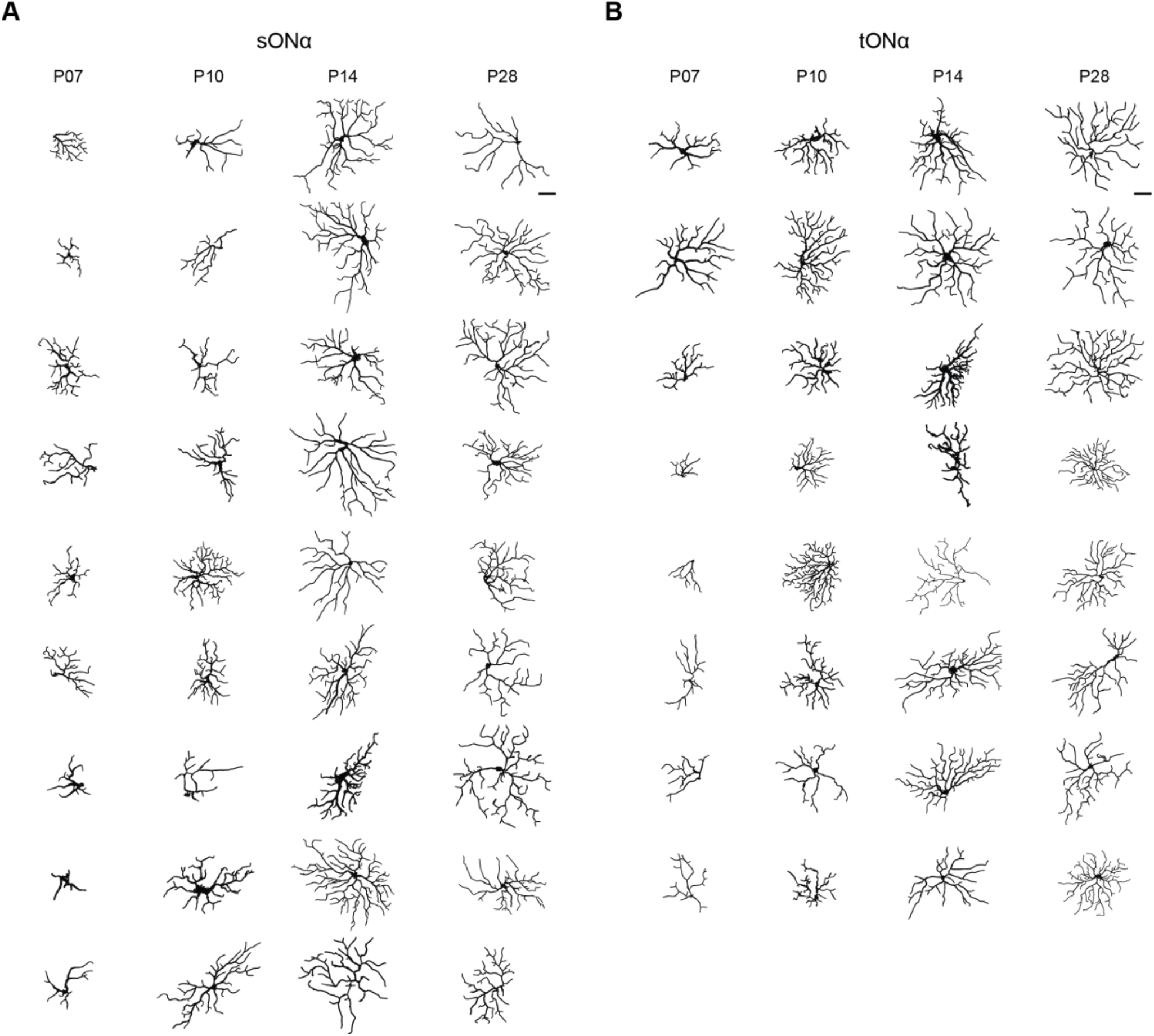
Representative dendritic reconstructions of sONα **(A)** and tONα **(B)** retinal ganglion cells at four different postnatal time points: P7, P10, P14, and P28. Each column represents cells from the same developmental stage, showing the morphological differences in dendritic arborization over time. The scale bar is 20 μm.

**Supplementary Figure 4.**
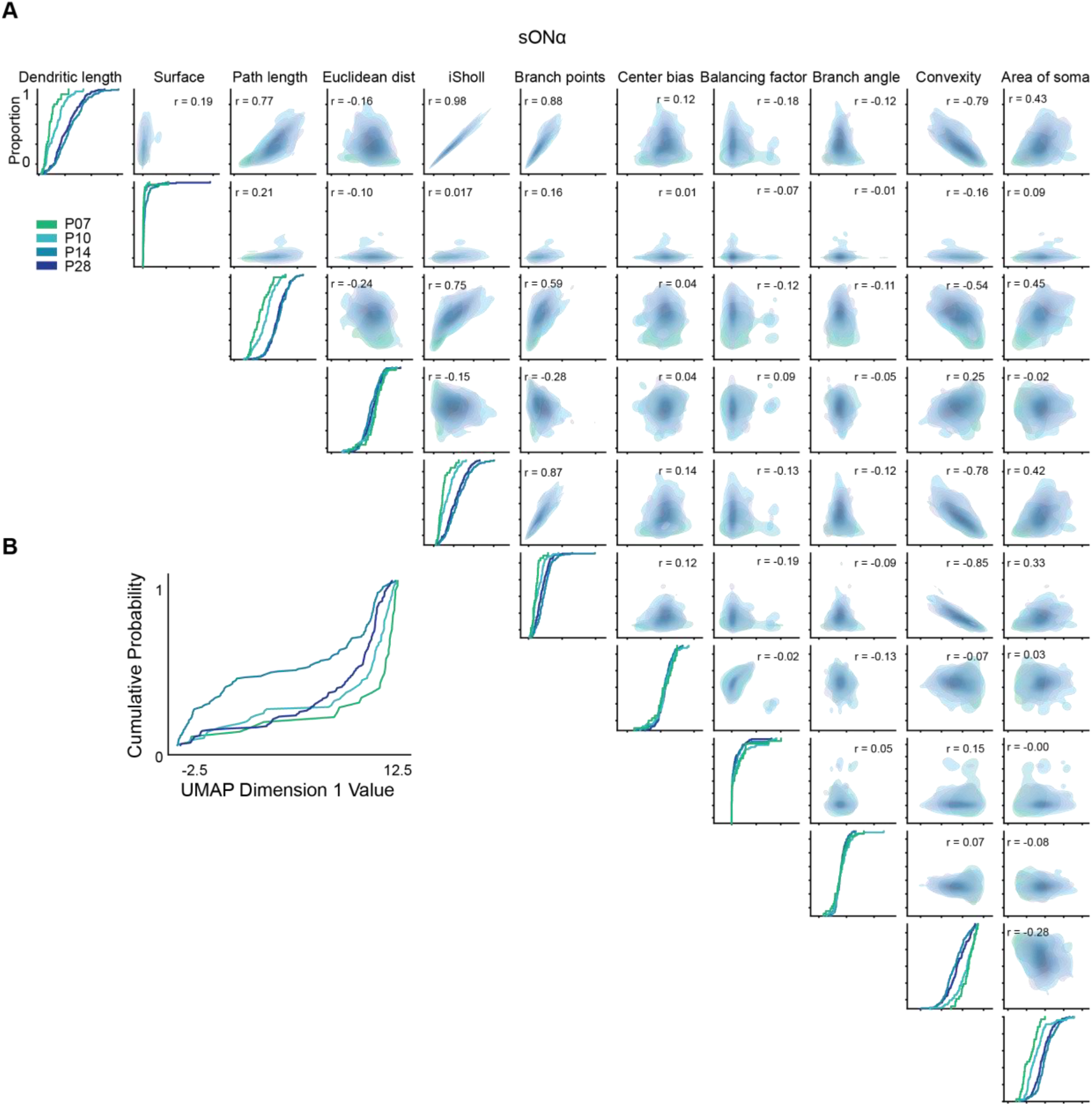
**(A)** Pairwise comparison of sONα cell parameters. Pairwise correlation analysis of key morphological parameters of sONα cells across developmental time points. Kernel density estimation (KDE) plots (blue shading) in the upper triangle illustrate the distribution of data points for each parameter combination. The diagonal panels display empirical cumulative distribution functions (ECDFs) for each parameter across different developmental stages. Correlation coefficients (r-values) are annotated in the upper triangle, quantifying pairwise relationships between parameters. **(B)** Cumulative distribution of UMAP Dimension 1. Cumulative probability distributions of Uniform Manifold Approximation and Projection (UMAP) Dimension 1 across developmental stages (P7, P10, P14, and P28) demonstrate the progressive structural differentiation of sONα cells over time. Each curve represents a different developmental time point, showing how the distribution of UMAP values evolves.

**Supplementary Figure 5.**
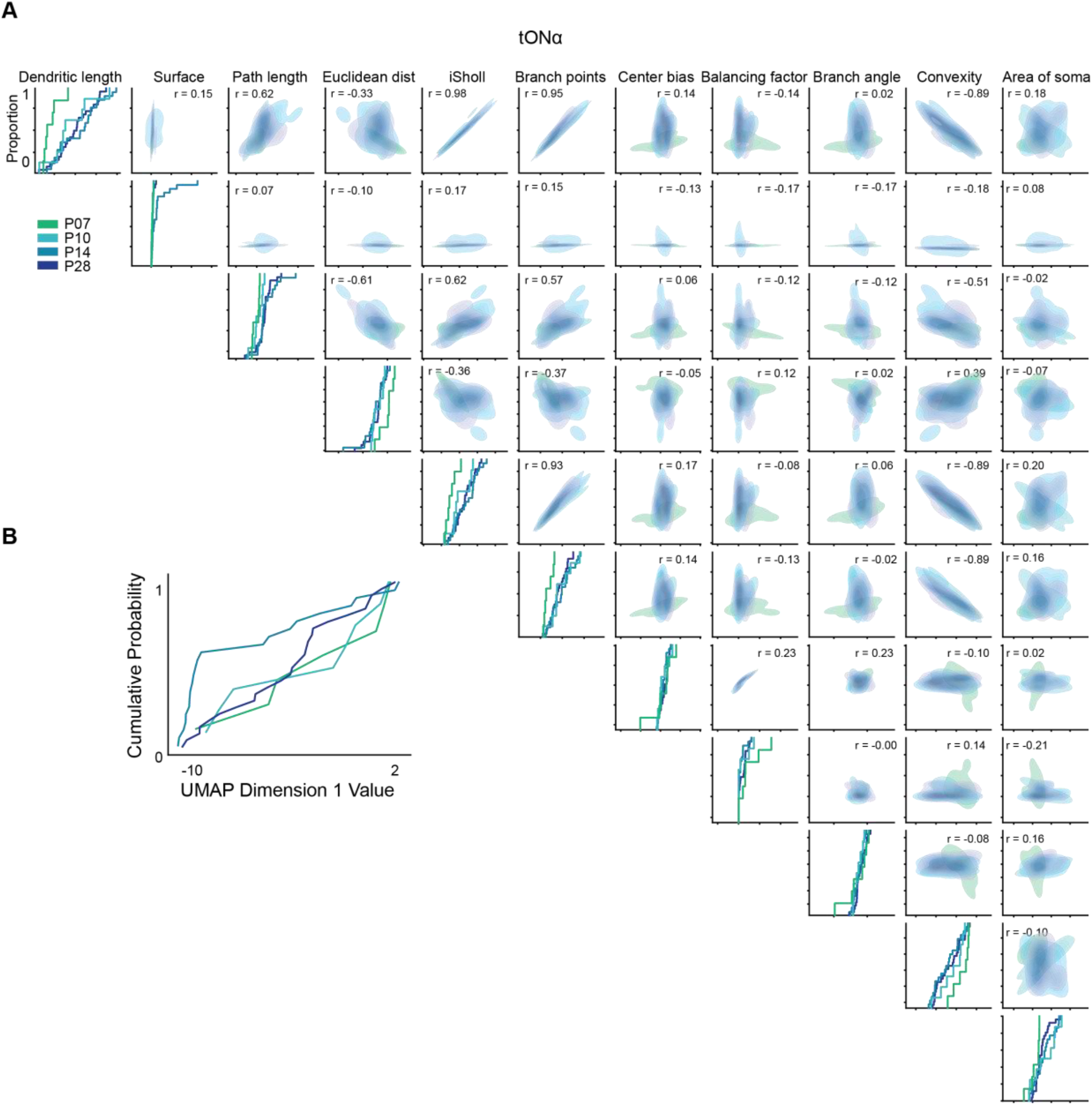
**(A)** Pairwise comparison of tONα cell parameters. Pairwise correlation analysis of morphological parameters across developmental time points. KDE plots in the upper triangle illustrate parameter distributions, while ECDFs are shown along the diagonal. Correlation coefficients (r-values) quantify pairwise relationships. **(B)** Cumulative distribution of UMAP Dimension 1. Cumulative probability distributions highlight progressive morphological differentiation across developmental stages.

**Supplementary Figure 6.**
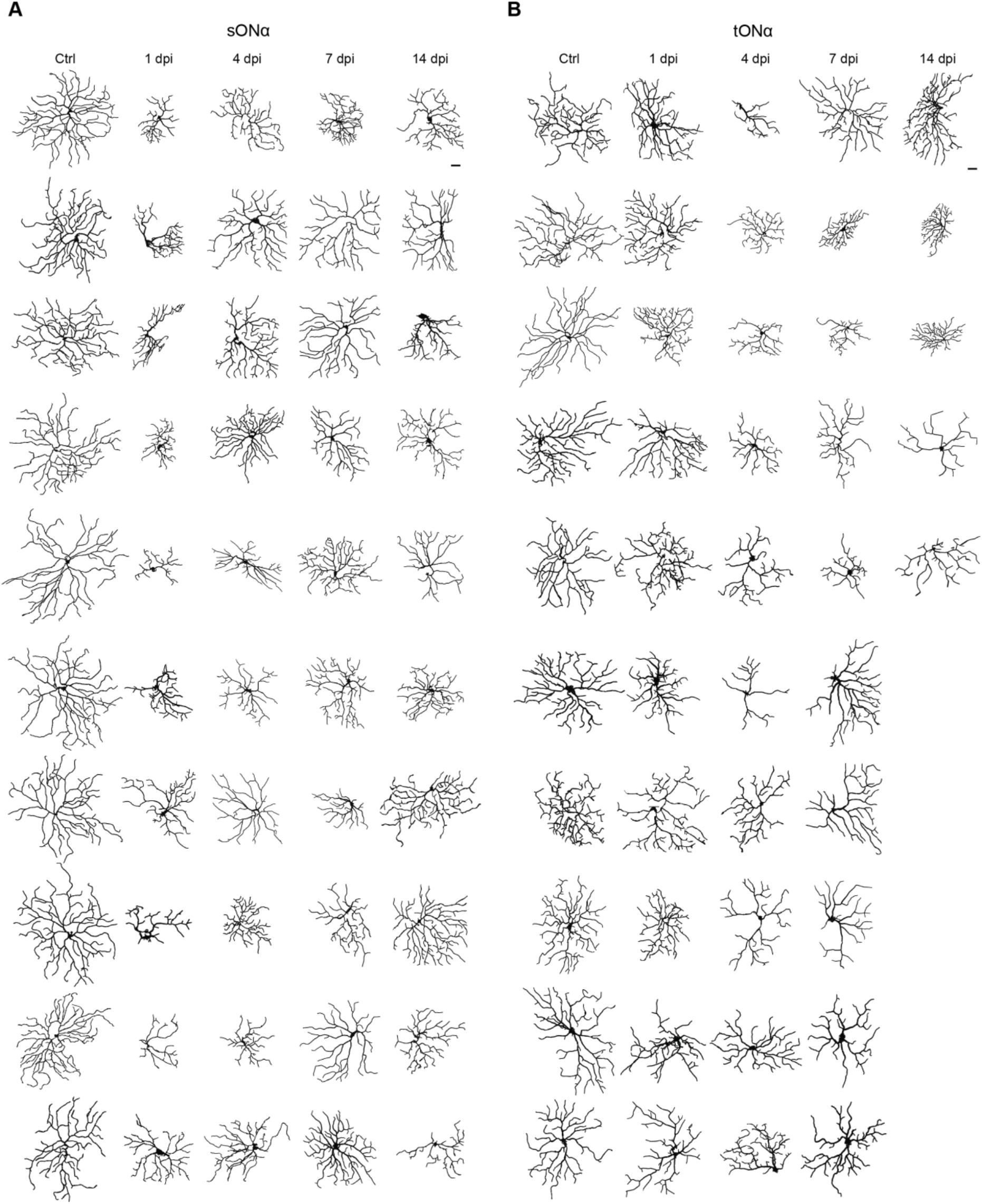
Example traces of sONα and tONα cells following optic nerve crush (ONC) at different time points. Representative dendritic reconstructions of sONα **(A)** and tONα **(B)** retinal ganglion cells after injury. Traces are grouped by Control (Ctrl) and days post-injury (dpi: 1, 4, 7, 14), showing morphological changes over time. Each column represents cells from the same condition, highlighting alterations in dendritic structure following ONC. Scale bar: 20 μm.

**Supplementary Figure 7.**
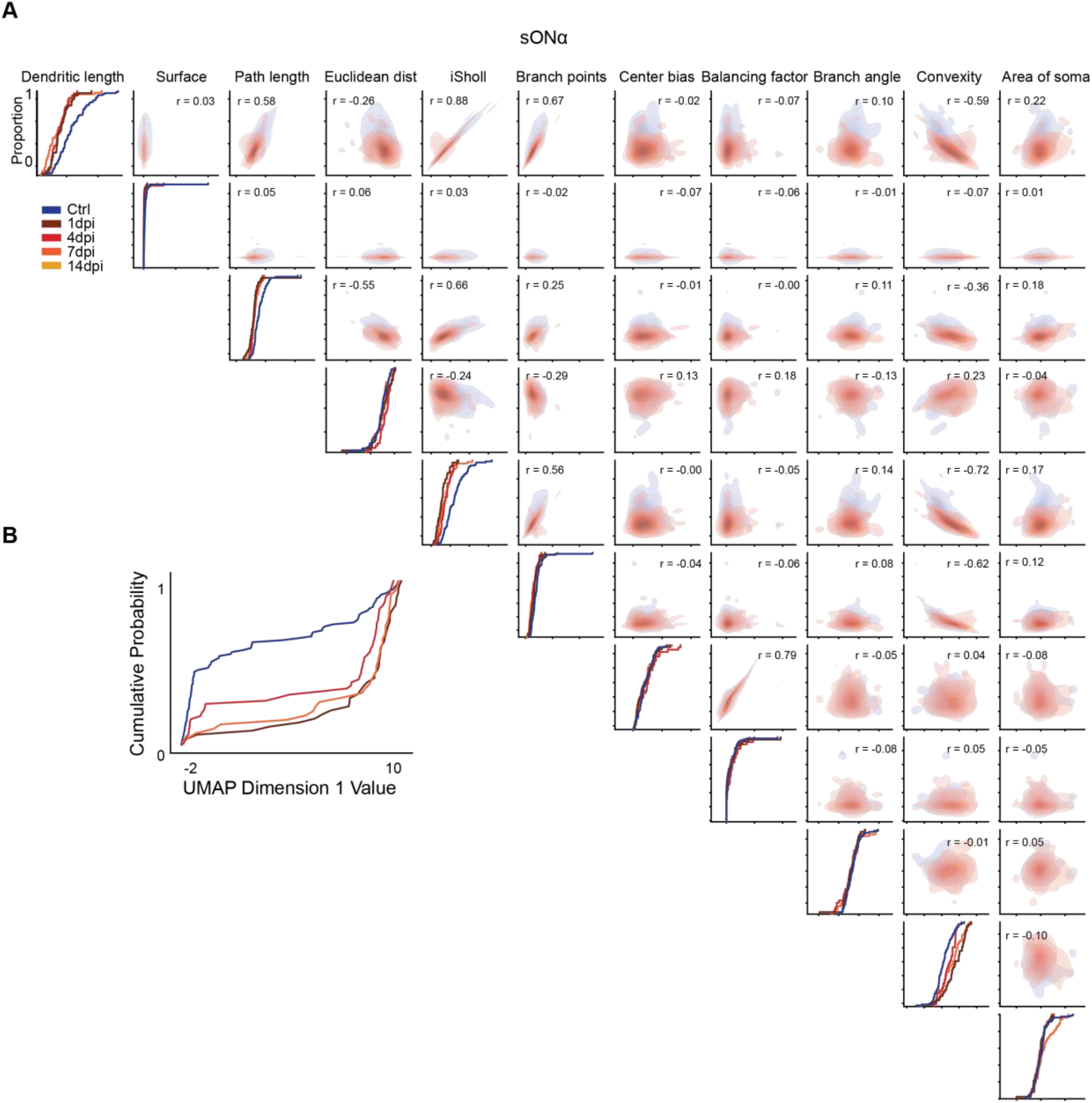
**(A)** Pairwise comparison of sONα cell parameters after injury. Pairwise correlation analysis of morphological parameters in sONα cells across different conditions: Control (Ctrl) and days post-injury (dpi: 1, 4, 7, 14). KDE plots (shaded in red) parameter distributions, while ECDFs in the diagonal panels show cumulative distributions. Correlation coefficients (r-values) are annotated in the upper triangle. **(B)** Cumulative distribution of UMAP Dimension 1. Cumulative probability distributions of UMAP Dimension 1 across control and post-injury time points, illustrating progressive morphological changes following ONC.

**Supplementary Figure 8.**
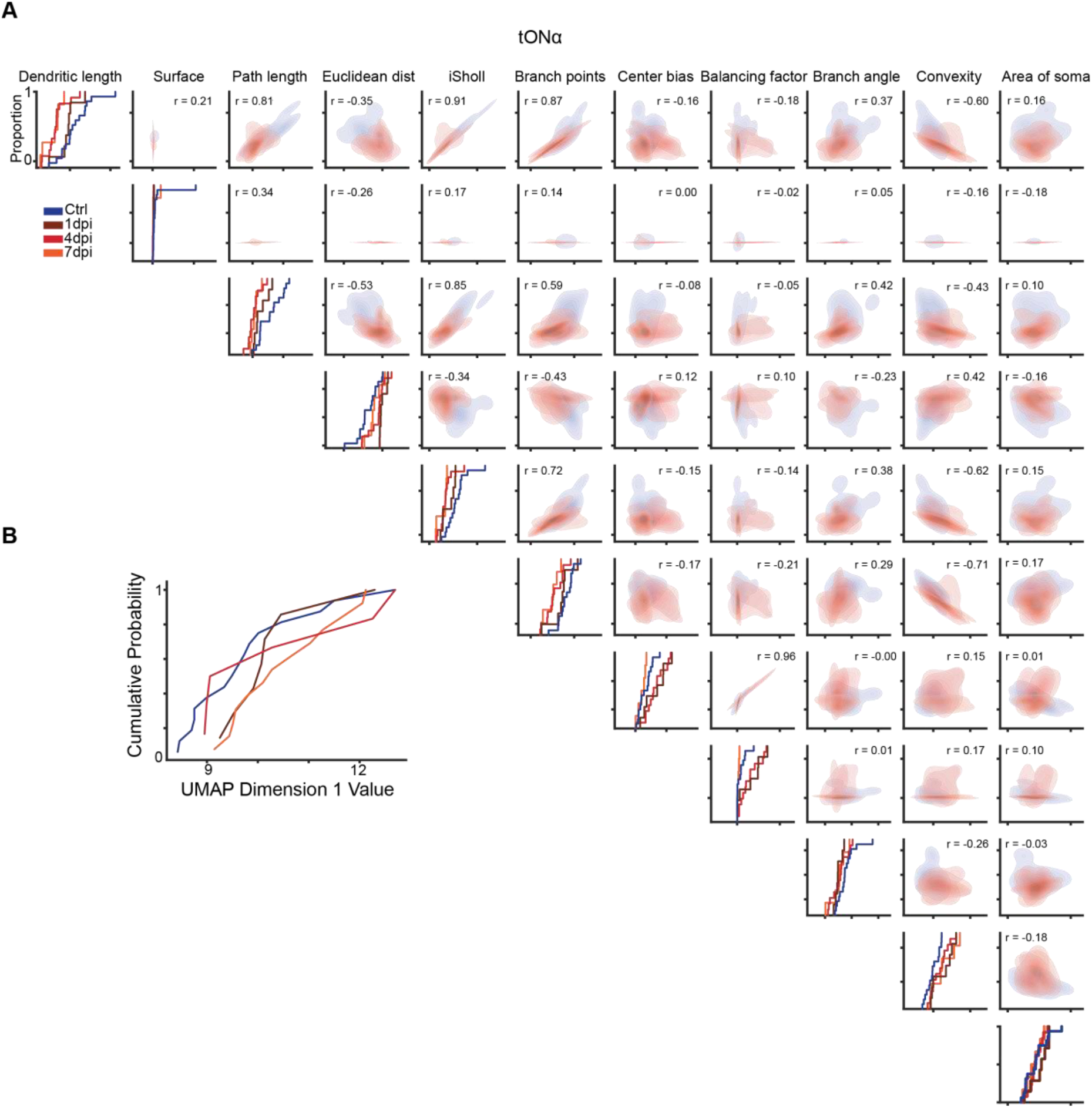
**(A)** Pairwise comparison of tONα cell parameters after injury. Pairwise correlation analysis of morphological parameters in tONα cells across different conditions: Control (Ctrl) and days post-injury (dpi: 1, 4, 7, 14). KDE plots (shaded in red) illustrate parameter distributions, while ECDFs in the diagonal panels show cumulative distributions. Correlation coefficients (r-values) are annotated in the upper triangle. **(B)** Cumulative distribution of UMAP Dimension 1. Cumulative probability distributions of UMAP Dimension 1 across control and post-injury time points, illustrating progressive morphological changes following ONC.

**Supplementary Figure 9.**
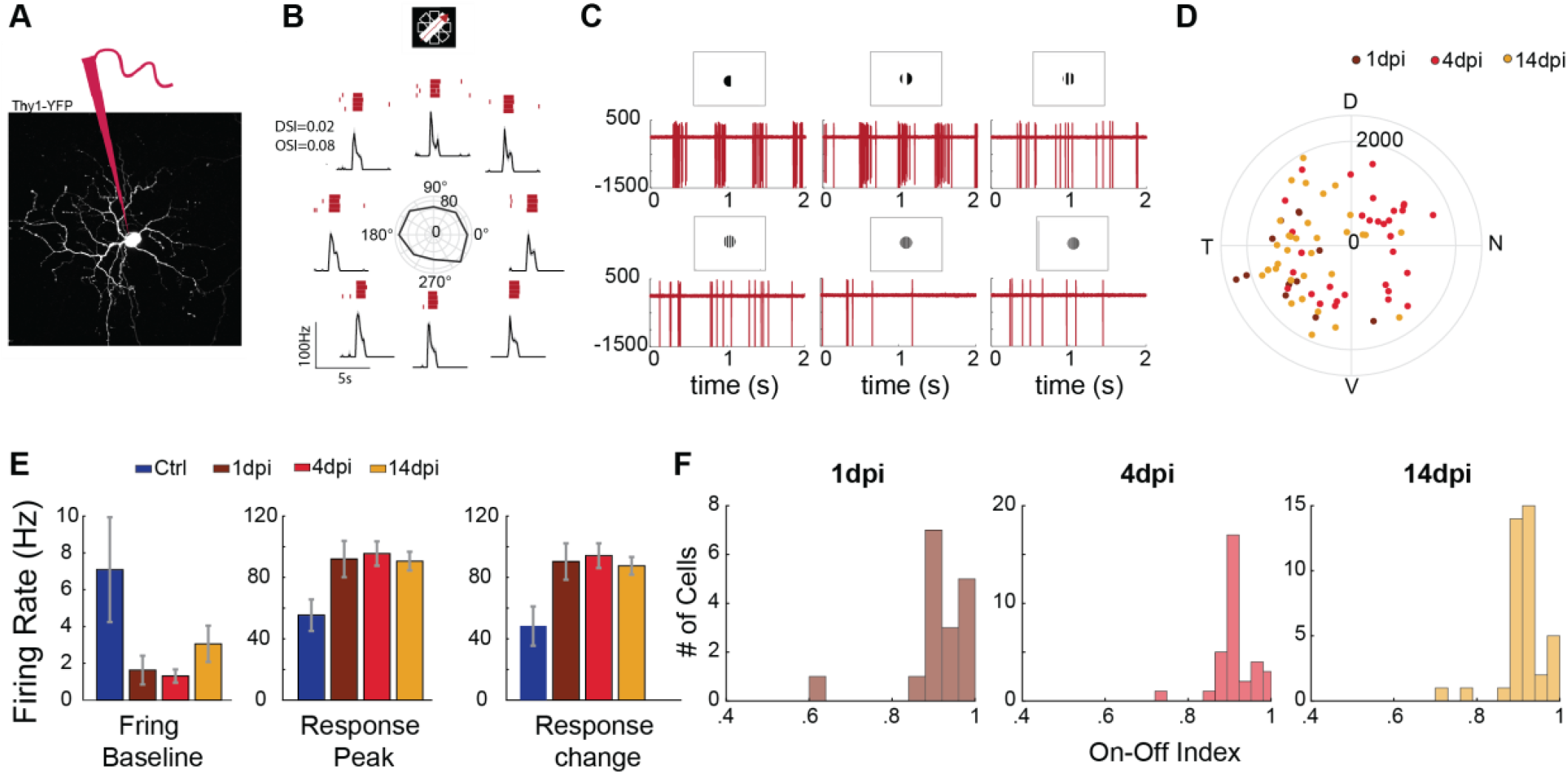
Targeted patch-clamp recording of Thy1-YFP retinal ganglion cells following optic nerve crush injury. **(A)** Maximum intensity projection of a two-photon image stack showing a Thy1-YFP-expressing cell following the optic nerve crush. **(B)** Example of the patched cell’s response to a moving bar. The traces (black) represent the mean firing rates across 10 trials, with individual raster plots shown above (red). Polar plot in the middle shows the broad tuning property. **(C)** Example firing spikes (red) showing responses to drifting gratings of varying spatial frequencies. **(D)** Spatial distribution of all patched ON alpha retinal ganglion cells across the retina. **(E)** Quantification of the patched ON alpha retinal ganglion cells’ activity: baseline firing rate before the response onset (left), peak response to the moving bar (middle), and the relative change in response (right). **(F)** On-Off index of the ON alpha cells at 1-, 4-, and 14-days post nerve crush.

**Supplementary Figure 10.**
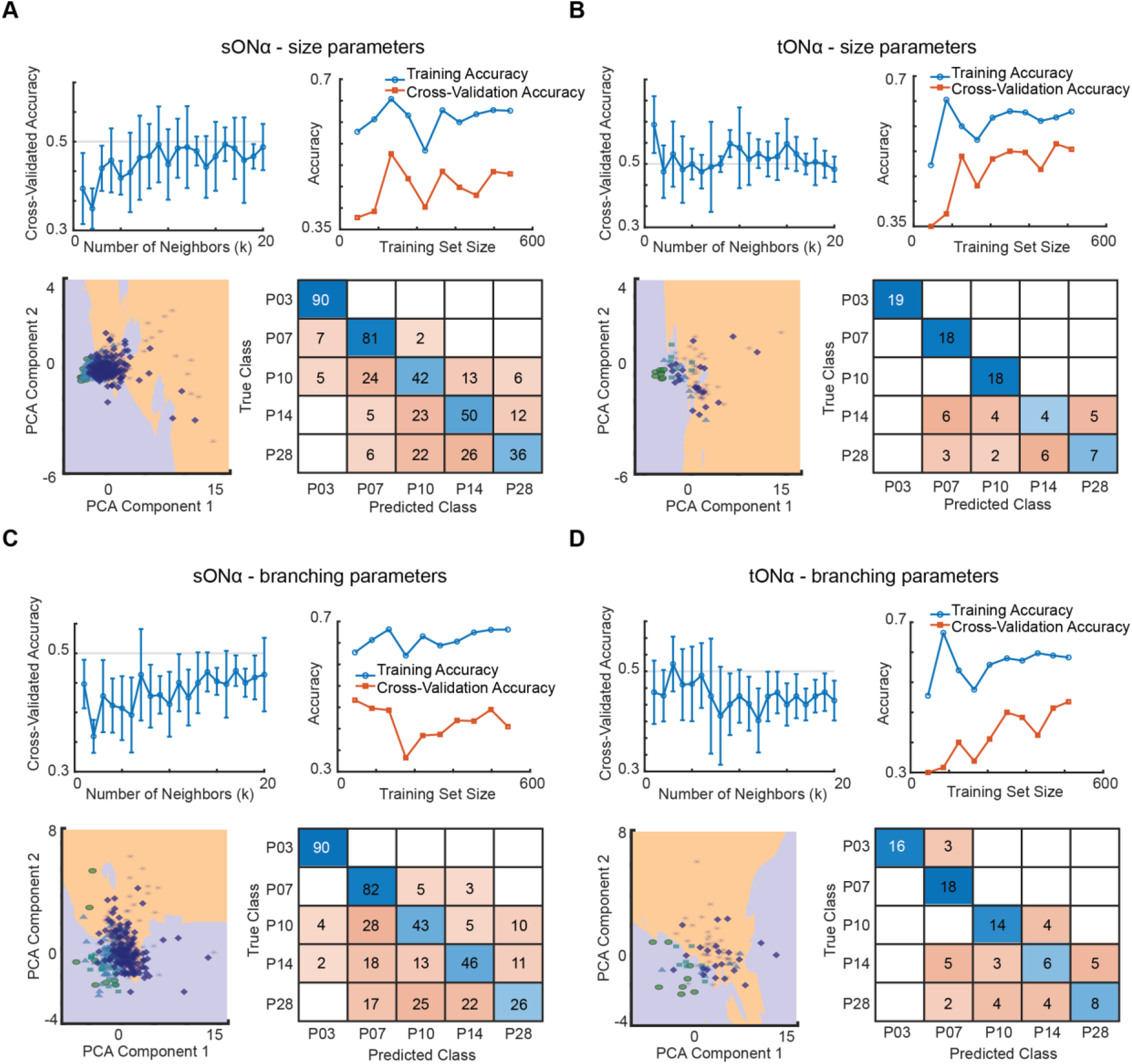
**(A)** sONα – Size features. i) k-NN Hyperparameter Tuning: The cross-validated accuracy is plotted against different values of k. Error bars indicate the standard deviation. ii) Learning Curve: Training accuracy (blue) and cross-validation accuracy (orange) are plotted against the number of training samples, showing model performance as more data is included. iii) Decision Boundary Visualization: The decision regions of the k-NN classifier are shown in the principal component (PCA) space for one selected value of k = 10. The scatter plot represents individual cells, color-coded by age group. iv) Confusion Matrix: The classifier’s performance in predicting cell age is visualized. Rows represent the true age category, while columns represent the predicted category. Correct classifications lie along the diagonal. **(B)** tONα – Size Features (B.i – B.iv) Same as (A), but for tONα cells using size-related features. **(C)** sONα – Branching Features (C.i – C.iv) Same as (A), but using branching-related features for sONα cells. **(D)** tONα – Branching Features (D.i – D.iv) Same as (C), but for tONα cells.

**Supplementary Figure 11.**
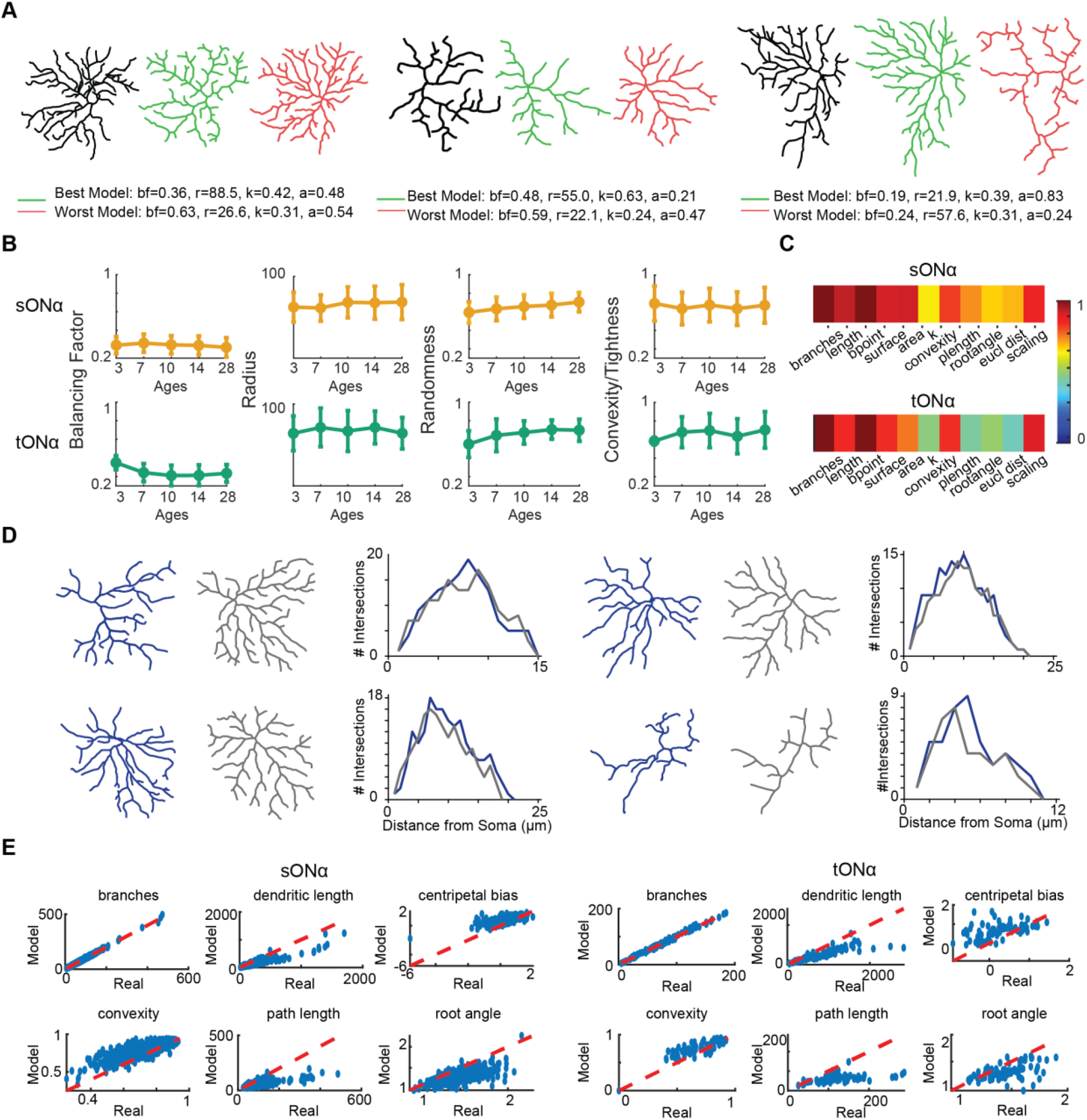
Comparison of real and modeled dendritic trees. **(A)** Representative examples of real (green) and modeled (red) dendritic structures. **(B)** Parameter optimization across development. Key parameters, including Balancing Factor, Radius, Randomness, and Convexity/Tightness, were optimized for different developmental stages (P03, P07, P10, P14, P28). Data points represent means, with error bars indicating standard deviation (SD). **(C)** Correlation heatmaps of parameter accuracy. Heatmaps display the Pearson correlation (r) between real and modeled values for key morphological parameters across developmental time points. Higher correlation values (yellow-red) indicate strong agreement between modeled and real data, while lower values (blue-green) suggest discrepancies. Correlation values range from 0 (no correlation) to 1 (perfect correlation). **(D)** Sholl analysis comparison between real and modeled trees. Sholl profiles were compared for real (gray) and modeled (blue) dendritic arbors, showing how well the model captures dendritic complexity at different developmental time points. **(E)** Scatter plots of real versus modelled data. Each subplot shows the relationship between real and modeled values for different morphological parameters. The red dashed line represents the identity line (perfect correlation), with points deviating from this line indicating discrepancies between modelled and real data.

